# Predicting resistance to synthetic lethal therapies in cancer

**DOI:** 10.64898/2026.10.12.757986

**Authors:** Metin Yazar, Christopher J. Lord, Colm J. Ryan

## Abstract

**Background:** Synthetic lethality (SL) occurs when an alteration in one gene renders cells vulnerable to inhibition of another, offering a strategy for targeted cancer therapy. PARP inhibitors represent the first clinically approved synthetic lethal therapy, and numerous additional candidates are now in clinical trials. Unfortunately, resistance to these therapies is common and impairs their overall effectiveness. Although there is a growing understanding of the factors that predict synthetic lethal interactions, the prediction of mechanisms driving resistance to these effects remains largely unstudied. Here, we sought to determine whether SL resistance could be predicted *a priori* using protein-protein interaction networks integrated with additional molecular profiles.

**Methods:** We curated genome-wide CRISPR-Cas9 chemogenetic resistance screens spanning ten unique SL biomarker-target pairs. For each pair, we extracted resistance genes and computed features capturing protein-protein interactions, gene expression, and essentiality. We compared these features between resistance and non-resistance genes and developed a random forest classifier to predict resistance. Model performance was assessed using random-split and structured leave-one-out schemes based on biomarkers, targets, and SL pairs to ensure robust generalisation. Independent validation of the model was performed by assessing its ability to predict resistance to two SL therapies currently in clinical trials, MTAP-PRMT5 and ATM-ATR.

**Results:** Resistance genes were significantly enriched among the protein-protein interaction partners of SL pairs across multiple datasets, suggesting functional interaction networks can aid the prioritisation of resistance mechanisms. Resistance genes tended to be more co-expressed with their associated target and also displayed distinct essentiality patterns. Although no individual feature was strongly predictive, integrating all features in a random forest model improved discrimination between resistance and non-resistance genes. Under leave-one-SL-pair-out cross-validation — our most stringent test of generalisation — the model achieved a mean area under the receiver operating characteristic curve (AUROC) of 0.72 across ten pairs (range 0.57-0.91; mean average precision ∼9%); a random train-test split gave a higher AUROC (∼0.78) that we treat as an optimistic upper bound, since biomarkers and targets are shared across folds. Average precision remained low in absolute terms, reflecting the inherent challenge of predicting extremely rare resistance events (<1% of genes).

**Conclusions:** Resistance to synthetic lethal therapies arises, at least in part, from predictable patterns of network connectivity, transcriptional coordination, and context-dependent essentiality. Our integrative framework provides a computational approach for anticipating resistance mechanisms across the expanding landscape of synthetic lethal therapies, offering a foundation for designing more durable and effective combination treatments.

## Introduction

Synthetic lethality (SL) enables selective killing of cancer cells with specific molecular alterations and can therefore be applied to uncover therapeutic targets in cancer (Gonçalves et al., 2025). The most well-known example in clinical use is the application of PARP inhibitors to treat select breast, ovarian, prostate and pancreatic cancer patients with defects in DNA repair by homologous recombination, including those with mutations in the *BRCA1* or *BRCA2* tumour suppressor genes (Bryant et al., 2005; Farmer et al., 2005). Multiple additional SL therapies are currently in different stages of clinical trials (Ahn et al., 2025; Burris et al., 2024; Okines et al., 2025; Simoneau et al., 2025), recently reviewed in (Gonçalves et al., 2025; Schäffer et al., 2024).

Functional genomics screens are an effective way to discover and model SL interactions in human cell lines (Dwane et al., 2021; McDonald et al., 2017; Shalem et al., 2014). Genetic loss-of-function screens systematically knock down or knock out genes to identify those whose loss is selectively lethal in a defined genetic background, thereby revealing SL partners. For example, large-scale screens revealed that cell lines with loss of *MTAP*, a gene that is often co-deleted with the tumour suppressor *CDKN2A*, display increased sensitivity to knockout or knockdown of protein arginine methyltransferase 5 (PRMT5) (Kryukov et al., 2016; Marjon et al., 2016; Mavrakis et al., 2016). Complementing these approaches, chemogenetic screens combine genome-wide CRISPR-Cas9 perturbation with drug treatment to identify genes whose loss confers sensitivity or resistance to a targeted therapy (Zhang et al., 2023) and form the basis of the resistance screens analysed in this study.

As with most targeted therapies, drug resistance to SL therapies is a major challenge and can reduce their durability and effectiveness (Huang et al., 2020; Li et al., 2020). Tumour heterogeneity and adaptive evolution enable cells to acquire or select for mutations that restore viability under therapeutic pressure. In SL contexts, such resistance may arise through changes in either of the synthetic lethal partners or via changes in other genes that re-establish the disrupted pathway or compensate for the inhibited target. For example, resistance to PARP inhibitors can occur via function-restoring reversion mutations in the biomarker gene *BRCA1* or via changes in the target gene *PARP1* (i.e. changes in the synthetic lethal partners) or via compensatory loss of members of the Shieldin complex (SHLD1, SHLD2, SHLD3 and REV7/MAD2L2) (Gupta et al., 2018; McCormick et al., 2017; Noordermeer et al., 2018). This complex acts downstream of 53BP1 to protect DNA double-strand-break ends from nucleolytic resection, thereby promoting non-homologous end joining at the expense of homologous recombination. In BRCA1-deficient cells, loss of Shieldin components restores end resection and homologous recombination, re-establishing the DNA-repair capacity that PARP inhibition is designed to exploit, and thereby drives resistance (Bunting et al., 2010; Dev et al., 2018; Jaspers et al., 2013; Noordermeer et al., 2018). Resistance can also arise through adaptive rewiring of signalling networks — for instance, reactivation of RAS-RAF-MEK-ERK signalling restores proliferative drive and confers resistance to targeted inhibitors of this pathway (Van Allen et al., 2014; Wagle et al., 2014). These examples suggest that drug resistance may arise not through mutation of random genes but through alteration of genes functionally related to the biomarker or target.

Protein-protein interaction (PPI) networks provide a powerful framework for understanding how genes act in concert to influence cellular fitness and drug response. PPIs have been widely used to predict and understand synthetic lethal (SL) interactions, as functionally related genes are more likely to exhibit synthetic lethality when perturbed (Benstead-Hume et al., 2019; Costanzo et al., 2016; De Kegel et al., 2021; Dey et al., 2024; Kelley & Ideker, 2005; Lord et al., 2020). Beyond pairwise interactions, it has been proposed that trigenic interactions play a significant role in regulating global biological processes, and also often occur among functionally related genes (Kuzmin et al., 2018). These highlight that polygenic relationships, rather than isolated gene effects, frequently underlie both synthetic lethality and therapeutic resistance in targeted therapies (Ryan et al., 2023).

There has been substantial work in both clinical and pre-clinical models to characterise mechanisms of resistance to PARP inhibitors (Wicks et al., 2022), but whether these mechanisms might indicate the presence of general principles regarding resistance that might apply in the context of other SL therapies remains completely unknown. In particular, it is unclear whether genes or pathways that might cause resistance can be predicted from prior knowledge. Being able to predict mechanisms of resistance *a priori* could have impact in terms of allowing the overall effectiveness of SL therapies to be improved or, at the very least, reduce the scale of efforts needed to identify mechanisms of resistance to a smaller number of candidates. Here, we hypothesise that resistance is not random, but instead follows patterns predictable from biological networks, in particular, genes that are closely connected to synthetic lethal partners in protein-protein interaction (PPI) networks may be more likely to drive resistance. We test whether network proximity, together with other molecular signals (essentiality and expression), can be integrated to predict resistance in a generalisable way. Our study has two main goals: (i) to test whether resistance genes are consistently enriched among the protein-protein interaction (PPI) partners of synthetic lethal (SL) pairs, and (ii) to turn this idea into a predictive machine learning model that remains reliable when tested on new biological settings and independent datasets.

## Results

### Resistance to PIK3CB inhibition in PTEN deficient models is driven by protein-protein interaction (PPI) partners of PTEN and PIK3CB

We hypothesised that resistance to synthetic lethal therapies might arise through alteration of genes known to have clear functional relationships with either the biomarker or target gene involved in the synthetic lethality. We initially sought to test this hypothesis on the well-established synthetic lethal relationship between the tumour suppressor *PTEN* (biomarker) and its synthetic lethal partner *PIK3CB* (target) (Wee et al., 2008). This synthetic lethality has been observed across cell lines from multiple cancer types, including breast and prostate cancers (Hancox et al., 2015; Wee et al., 2008), and appears to be among the most robust or penetrant synthetic lethal interactions reported to date (Lord et al., 2020). Nonetheless, not all *PTEN* deficient models are sensitive to PIK3CB inhibition (Iorio et al., 2016) and a recent CRISPR-Cas9 screen identified multiple genes whose loss conferred resistance to a small molecule PIK3CB inhibitor in *PTEN* deficient models (Dunn et al., 2022). This latter screen identified genes whose loss conferred resistance to the PIK3CB inhibitor AZD8186 in three different PTEN null breast cancer models, for a total of 99 genes that cause resistance in at least one cell line. Using this as a test case, we obtained the interaction partners for PTEN and PIK3CB from the STRING and BioGRID interaction databases (see methods) and analysed the overlap of these interactors with the resistance genes reported in the CRISPR screen (Fig. 1A). The BioGRID contains protein-protein interactions supported by direct experimental evidence, while STRING contains *functional* interactions including those derived from membership of the same annotated pathway. Many of the reported resistance genes had a previously reported interaction with PIK3CB (e.g. *SOCS6*), PTEN (e.g. *TSC1*, *STK11*), or both (e.g. *TSC2*, *PIK3R3*) (Fig. 1C). Indeed, we found that the resistance genes were significantly enriched among known interaction partners of PTEN and PIK3CB (Fig. 1B) (STRING: OR = 5.08, p = 1.71e-09; BioGRID physical: OR = 3.19, p = 1.35e-03, both with Fisher’s exact test). At least for the specific case of PTEN-PIK3CB, this supports the hypothesis that resistance to synthetic lethality is enriched among the protein-protein interaction partners of the synthetic lethal pair.

**Fig 1.**
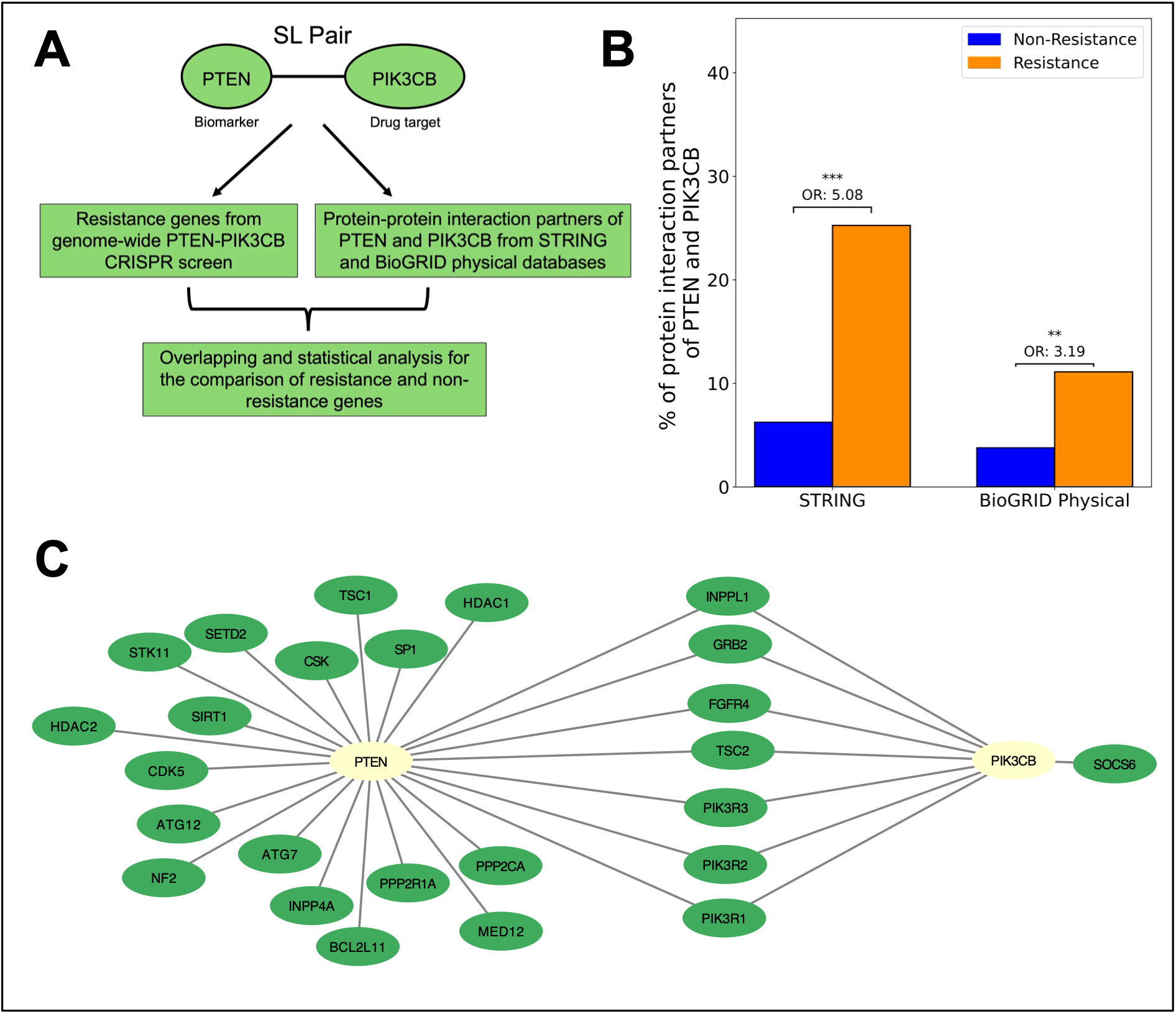
PTEN-PIK3CB resistance genes are enriched among the protein-protein interaction (PPI) partners of PTEN and PIK3CB. **A)** Workflow for the comparison of the resistance and non-resistance genes among PPI partners of PTEN (biomarker) and PIK3CB (target). **B)** Bar charts showing, for resistance (orange) and non-resistance (blue) genes, what percentage of them have an interaction with either PTEN, PIK3CB or both in the BioGRID physical and STRING (medium-confidence network, interaction score ≥0.4) databases (*** for p < 0.001, ** for p < 0.01, * for p < 0.05). **C)** Network diagram (visualised with Cytoscape (3.10.2) (Shannon et al., 2003)) showing the STRING interactions between PTEN, PIK3CB and the genes identified as causing resistance in the CRISPR screen.

### Resistance is enriched among PPI partners of multiple SL pairs

To assess whether our findings from PTEN-PIK3CB extended to the resistance mechanisms of other synthetic lethal gene pairs, we obtained the results of genome-wide chemogenetic CRISPR-Cas9 resistance screens for additional targeted therapies. We defined synthetic lethality broadly as interactions where a biomarker (mutated gene e.g. *PTEN*) was associated with increased sensitivity to inhibition of a target (e.g. PIK3CB) and the biomarker was not the same gene as the target. This led to the inclusion of resistance screens for synthetic lethal interactions involving oncogenes (e.g. NRAS-CDK4/6) (Hayes et al., 2019) but not screens for resistance to therapies that exploit oncogene addiction (e.g. to assess resistance to BRAF inhibitors in *BRAF* mutant models). Resistance genes were defined as those whose gene loss via CRISPR-mediated knockout conferred resistance to the SL therapy. In total, we collected the results for ten unique synthetic lethal combinations across nine different publications. Some SL combinations were screened multiple times in different publications (e.g. *KRAS-MEK1/2*, *BRCA1-PARP1, NRAS-MEK1/2*) and some were screened multiple times within a single publication in distinct cell lines, with distinct inhibitors of the same target, or in independent clonal derivatives of one line (Table 1). To avoid duplication, the results from all screens targeting the same gene pair were aggregated together — both within studies, when multiple cell lines were used such as the 3 cell lines used to study *PTEN*-*PIK3CB* resistance, and across studies, when the same gene pair was studied independently in multiple publications (e.g. *BRCA1-PARP1* was screened in Dev et al., 2018 and Noordermeer et al., 2018). Similarly, results exploiting the same SL using distinct inhibitors (e.g. the PARP inhibitors olaparib and talazoparib, both exploiting the *BRCA1*-PARP1 SL) were aggregated together. For aggregated results, resistance labels were defined by a union rule: a query gene was considered ‘Resistance’ if labelled as a resistance gene in *any* included screen for that SL pair. Most screens identified between 8 and 214 resistance genes (median 100), suggesting that genes whose loss causes resistance to a given SL are extremely rare (∼0.5% of the screened library). In general, the hits defined by different publications identifying resistance to the same SL pair had a moderate, but highly significant, overlap (Fig. S1).

**Table 1:** List of published genome-wide CRISPR-Cas9 chemogenetic drug resistance screens analysed.

| **Biomarker** | **Drug target** | **Cell line** | **Drug** | **Author** |
| --- | --- | --- | --- | --- |
| *PTEN* | *PIK3CB* | HCC-70, EVSA-T, ZR-75-1 | AZD8186 | (Dunn et al., 2022) |
| *PTEN* | *AKT* | HCC-70, EVSA-T, ZR-75-1 | capivasertib | (Dunn et al., 2022) |
| *ARID1A* | *ATR* | YCC6 | berzosertib | (Llorca-Cardenosa et al., 2022) |
| *BRCA1* | *PARP1* | SUM149PT | olaparib, talazoparib, AZD2461 | (Dev et al., 2018) |
| *BRCA1* | *PARP1* | SUM149PT, BRCA1-KO RPE1 | olaparib, talazoparib | (Noordermeer et al., 2018) |
| *BRCA2* | *PARP1* | BRCA2-knockout HeLa | olaparib | (Clements et al., 2020) |
| *CCNE1* | *PKMYT1* | FT282-hTERT | RP-6306 | (Gallo et al., 2022) |
| *KRAS* | *MEK1/2* | PATU8902, PATU8988T | trametinib | (Wang et al., 2017) |
| *NRAS* | *MEK1/2* | MELJUSO | trametinib | (Hayes et al., 2019) |
| *NRAS* | *CDK4/6* | MELJUSO | palbociclib | (Hayes et al., 2019) |
| *BRAF* | *MEK1/2* | HCC364 | trametinib | (Krall et al., 2017) |
| *KRAS* | *MEK1/2* | CALU1 | trametinib | (Krall et al., 2017) |
| *NRAS* | *MEK1/2* | H1299 | trametinib | (Krall et al., 2017) |

Analysing the overlap of biomarker/target PPI partners and the resistance *vs.* non-resistance genes in these screens showed that the enrichment of the partners was statistically significant in 9 of 10 unique SL pairs using the STRING interaction network (Fig. 2A) and 6 of 10 SL pairs with the BioGRID network (Fig. S2). Rather than just assessing each screen individually, we also assessed the enrichment of PPI partners among the resistance and non-resistance genes of all the screens combined (Fig. 2B) (STRING: OR = 3.34, p = 5.82e-49; BioGRID physical: OR = 2.41, p = 7.44e-23, See methods). Furthermore we found that this enrichment was still evident when only analysing SL pairs with a tumour suppressor as a biomarker (STRING: OR = 3.87, p = 4.29e-34; BioGRID physical: OR = 3.02, p = 2.57e-20 both with Fisher’s exact test), when excluding the PARP inhibitor screens (STRING: OR = 3.30, p = 1.18e-38; BioGRID physical: OR = 2.34, p = 7.56e-15 both with Fisher’s exact test), and when excluding screens involving MEK inhibitors (the most widely studied drug in our collection) (STRING: OR = 4.12, p = 2.11e-45; BioGRID physical: OR = 3.22, p = 1.65e-26 both with Fisher’s exact test) (Fig. S3). This suggested that the enrichment of resistance genes among the protein-protein interaction partners of SL genes is not an artefact of any one biomarker, target, or SL pair.

**Fig 2.**
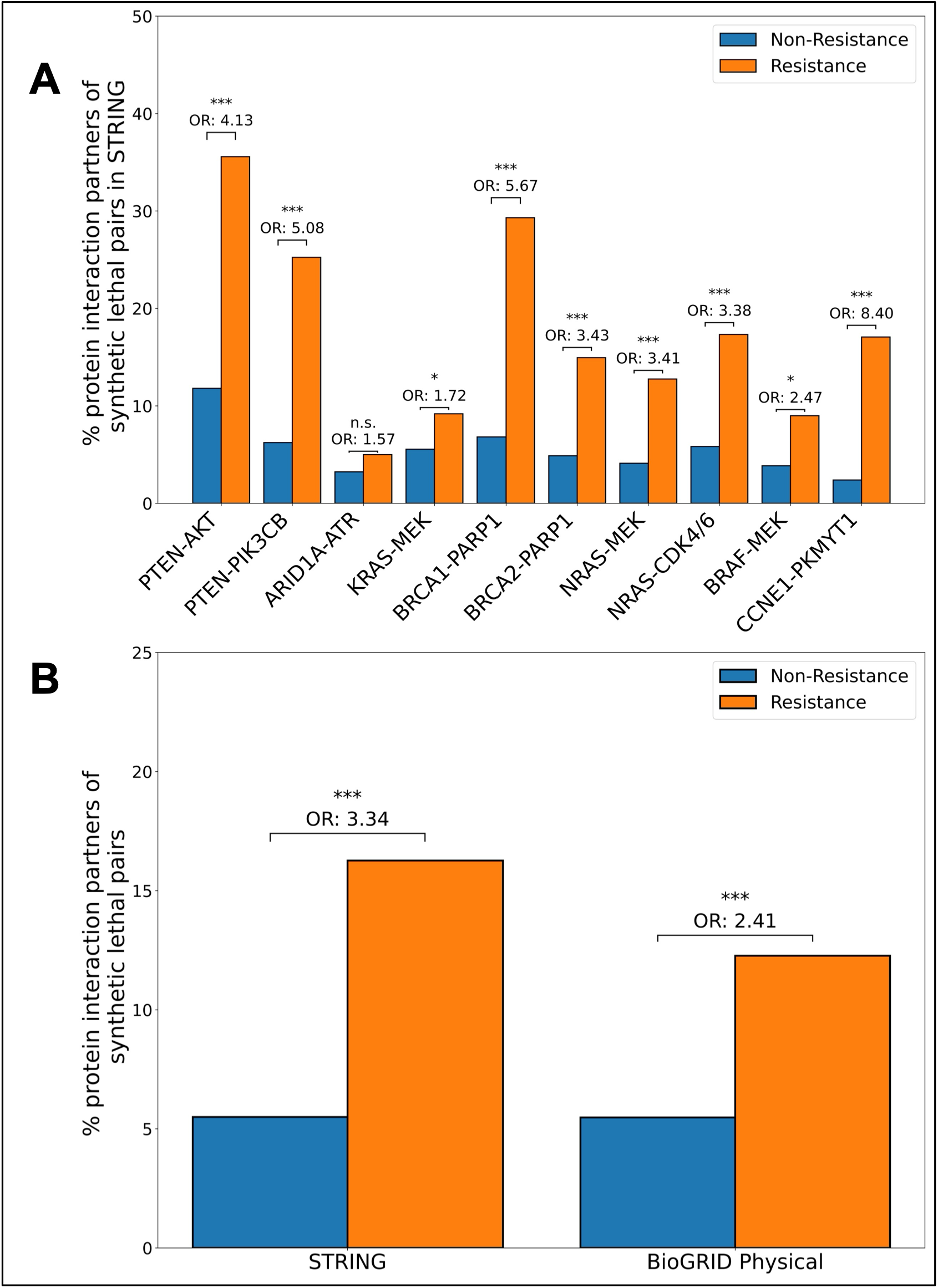
Resistance genes are overrepresented in protein-protein interaction (PPI) partners of multiple synthetic lethal pairs. **A)** Percentages of resistance (orange) and non-resistance (blue) genes that overlap with known PPI partners of each synthetic lethal (SL) pair in the STRING database. Statistical enrichments were calculated using Fisher’s exact test, reporting odds ratios and p-values (***p < 0.001, **p < 0.01, *p < 0.05, n.s. = not significant). **B)** Aggregated percentages of resistance (orange) and non-resistance (blue) genes overlapping with PPI partners across all genome-wide CRISPR-Cas9 chemogenetic screens, shown for both STRING and BioGRID physical databases (***p < 0.001, **p < 0.01, *p < 0.05, n.s. = not significant).

Previous work has established that ascertainment bias can result in some genes, especially known cancer drivers, displaying artefactually high numbers of connections on aggregated interaction networks such as those from STRING and BioGRID (Rolland et al., 2014). When analysing individual SL pairs this was not a significant concern, as we compared the observed connectivity of the resistance genes with that expected based on the degree of their associated SL pair. However, when analysing patterns of connectivity for the resistance genes across all SL gene pairs combined, this had the potential to impact our results. For this reason, we also compared the observed connectivity of ‘resistance’ genes with 1,000 degree-matched randomisations of both STRING and BioGRID physical networks and counted how many edges there were in common each time, to build up an ‘expected overlap’ distribution. For both BioGRID and STRING, we observed significantly more interactions between resistance genes and the biomarker/target than would be expected by chance (p < 0.05 and p<0.01 respectively) (Fig. S4). These results demonstrated that resistance genes cluster non-randomly around SL partners in PPI space, reinforcing that network proximity is a defining feature of resistance mechanisms and not simply a consequence of node degree. The fact that STRING, which incorporates both protein-protein interactions and broader functional relationships such as membership of the same pathway, shows a clearer signal than BioGRID suggests that it is not just physical protein-protein interactions that drive the signal.

### Gene essentiality and expression derived features are predictive of SL resistance

Our analyses to this point indicate that PPI and functional interaction networks can contribute to the identification of candidate resistance genes. Nevertheless, most established PPI partners of synthetic lethal pairs do not confer resistance; conversely, not all resistance genes have an established PPI with the biomarker or target. To address these limitations, we sought to determine whether additional molecular features could improve resistance prediction in synthetic lethal contexts. We calculated seventeen features describing both the potential resistance genes and their relationship to the biomarker or target in question (Fig. 3).

**Fig 3.**
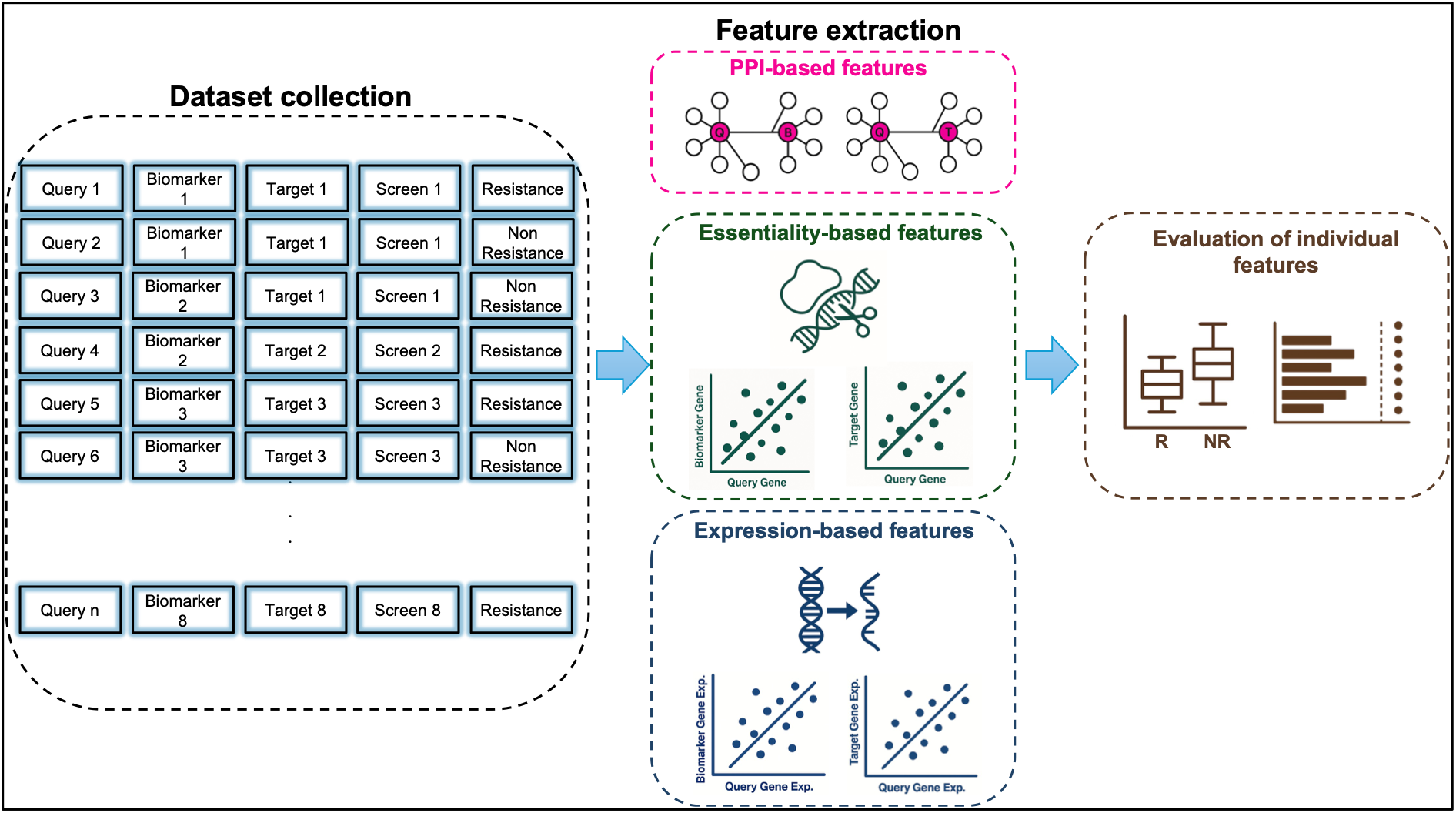
Overview workflow of data preprocessing for predicting drug resistance to SL therapies. This figure provides a schematic overview of the workflow used for data preprocessing, feature extraction and evaluation of the features. The dataset collection (left, blue box) integrates query genes from genome-wide CRISPR resistance screens, annotated with biomarker, target, screen identifiers, and resistance labels. Feature extraction (middle) is grouped into three main categories: protein-protein interaction (PPI)-based features (pink box), which quantify direct and shared protein-protein interactors between query genes and synthetic lethal partners; essentiality-based features (green box), derived from genome-scale CRISPR dependency data to capture functional relationships between query genes and biomarkers/targets; and expression-based features (blue box), which incorporate co-expression patterns and baseline transcriptional properties across tissues. In the final stage (feature evaluation, brown box), each feature was assessed individually for its ability to distinguish resistance from non-resistance genes using performance metrics such as area under the ROC curve (AUC), average precision, and statistical significance (Mann-Whitney U test).

The features are predominantly quantitative and could be grouped into three broad categories, expression, essentiality and protein-protein interaction based (Supplementary File-1). Expression-based features contain the average and variance of mRNA expression of the query gene in the Genotype-Tissue Expression (GTEx) database (GTEx Consortium, 2020) and of the expression correlation between query-biomarker and query-target. Features in the protein-protein interaction category derived from the interaction score between the query-biomarker and the query-target in STRING (Szklarczyk et al., 2023) and their interaction status in BioGRID Physical (Oughtred et al., 2021) as well as from the number of shared protein-protein interactors in the STRING medium confidence network and the BioGRID physical network (computed as the signed –log10 p value from a Fisher’s exact test of the overlap of query and biomarker/target interactors) (Oughtred et al., 2021). We calculated the essentiality-based features from DepMap gene effect scores (Meyers et al., 2017); these include the co-essentiality correlation between query-biomarker and query-target pairs, the variance and the average essentiality score of the query gene across cell lines, and the proportion of cell lines in which the query gene is essential.

To assess each feature’s ability to distinguish non-resistance (n = 179,591) from resistance (n = 1,440) gene triplets (query-biomarker-target) in the dataset, we treated each of the seventeen query-level features as a standalone classifier and evaluated it using standard machine learning metrics such as area under the receiver operating characteristic curve (AUROC) and average precision (AP). We also applied a statistical test (Mann-Whitney U test) to the fifteen continuous features among them (Fig. 4 and Fig. S5). The best predictor of resistance in our dataset is the query gene essentiality variance (∼0.73, average precision ∼2.20%), followed by shared protein-protein interactors of query-target in BioGRID (∼0.68, average precision ∼1.91%), essentiality percentage per cell line (∼0.68, average precision ∼1.43%) and co-expression between query and target gene (∼0.67, average precision ∼1.41%). It should be emphasised that the baseline average precision by chance is approximately 0.8%, reflecting the substantial class imbalance. This implies that, given the imbalance, even well performing classifiers will naturally exhibit relatively low average precision scores when viewed in absolute terms.

**Fig 4.**
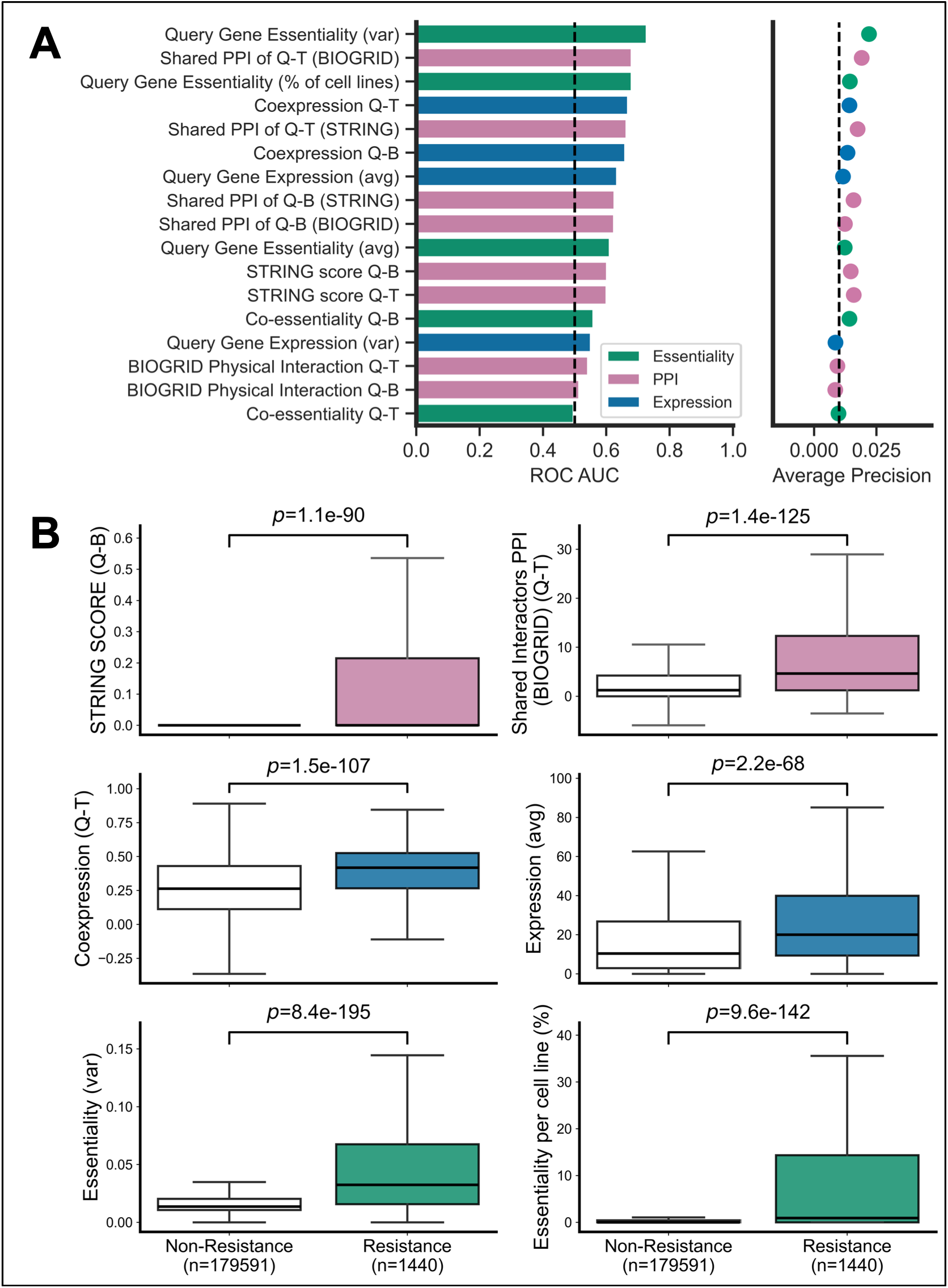
Resistance genes display distinct network, expression, and essentiality profiles. **A**) Receiver operating characteristic (ROC) and average precision (AP) metrics for individual features treated as standalone classifiers. Features span three major categories: protein-protein interaction (PPI) (pink), expression (blue), and essentiality (green). Bars represent AUROC scores, and corresponding AP values are shown as coloured dots, with feature categories indicated by colour. **B)** Representative boxplots showing the distributions of selected features between resistance and non-resistance genes. Comparisons include STRING interaction score between query and biomarker (Q-B), co-expression between query and target (Q-T), query gene expression (average), shared interactors in BioGRID (Q-T), query gene essentiality variance, and percentage of cell lines in which the query gene is essential. Statistical significance of group differences was assessed with two-sided Mann-Whitney U tests, with p-values indicated above each comparison.

Consistent with the AUROC analysis, most of the features (14 out of 15 continuous features) had statistically significant differences in distribution between resistance and non-resistance genes (Mann-Whitney U test p < 0.001) (Fig. 4B and Fig. S5). Resistance genes have higher STRING protein-protein interaction scores with both their SL pairs (query-biomarker (p = 1.1e-90) and query-target (p = 4.0e-96)), and greater shared physical interactors in BioGRID (query-biomarker (p = 5.6e-61) and query-target (p = 1.4e-125)) and shared STRING medium confidence network (query-biomarker (p = 1.2e-61) and query-target (p = 1.0e-104)), suggesting closer functional proximity within biological networks. These differences are consistent with pathway-level mechanisms of SL therapy in cancer, where redundant or compensatory pathways buffer single-node inhibition (Holohan et al., 2013). Resistance genes also demonstrated elevated average expression (p = 2.2e-68) and higher co-expression levels (query-biomarker (p = 8.8e-97) and query-target (p = 1.5e-107)). These results suggest that genes whose loss causes resistance tend to be more highly expressed and may be transcriptionally co-regulated with either the biomarker or target gene.

Notably, essentiality variance (p = 8.4e-195) emerged as the most discriminative feature between resistance and non-resistance genes and as noted had the highest AUROC and AP. The essentiality percentage, defined as the proportion of cell lines in which a query gene is classified as essential in DepMap using a predefined gene effect threshold (gene effect score < –0.6), was also significantly different in resistance genes compared to non-resistance genes (p = 9.6e-142). These findings suggest that genes whose loss causes resistance to some synthetic lethal therapies (i.e. restore viability) are more likely than other genes to cause context-specific fitness defects. Previous studies of PARP inhibitor resistance in BRCA1-deficient cells have identified multiple components of the 53BP1-Shieldin pathway whose loss restores homologous recombination and drives resistance. These include 53BP1-RIF1 axis members such as TP53BP1, as well as Shieldin complex components SHLD1, SHLD2, SHLD3 and MAD2L2*/*REV7 (Noordermeer et al., 2018). Importantly, loss of these same factors has also been shown to confer hypersensitivity to POLQ inhibition, establishing a well-characterised double-bind whereby restoration of homologous recombination through Shieldin or 53BP1 loss creates a new dependency on POLQ-mediated repair (Zatreanu et al., 2021). Consistent with our general findings, these factors show elevated essentiality variance relative to the genome-wide background (e.g. *REV7* = 0.17 and *TP53BP1* = 0.05, versus a non-resistance median of 0.01), reflecting context-specific essentiality. Consistent with higher variance, these genes are also essential in more cell lines – *REV7*, for example, is classified as essential in ∼73% of DepMap cell lines, far above the median gene.

Collectively, these analyses demonstrate that resistance in synthetic lethal contexts is not driven by a single dominant molecular characteristic, but is associated with multiple molecular properties, including protein-protein interaction connectivity, expression patterns, and essentiality.

### An integrative random forest model leveraging distinct molecular features improves prediction of resistance across diverse synthetic lethal contexts

Since multiple features were identified as individually predictive of drug resistance in cancer, we hypothesised that integrating these features might be even more predictive. Therefore, we integrated all features using a random forest based classifier (Breiman, 2001) as a predictive model for SL resistance. In addition to the seventeen features evaluated above, we also incorporated a binary feature indicating whether the biomarker was subject to an activating or loss-of-function mutation, to distinguish between patterns that might be different for oncogenes and tumour suppressors. Initially, we evaluated our classifier by performing training-test split with triplets (biomarker, target, query) allocated at random to the train or test set. As a benchmark we compared against the STRING score (between biomarker and query), to assess whether the integrated classifier outperformed the STRING score alone (Fig. 5A). Our model achieves a mean AUROC of 0.78 ± 0.02 averaged across 5 folds and average precision ∼10% on the test data, while STRING achieved an AUROC of ∼0.60 and average precision of ∼1%. However, random splits in this setting can overestimate performance because observations are not independent: the same biomarker may be screened against multiple distinct targets (e.g. *PTEN* is screened against both PIK3CB and AKT inhibitors) while the same target may be screened against multiple biomarkers (e.g. MEK inhibitors are screened in the context of *KRAS* and *BRAF* mutation). This can enable data leakage (biomarker or target specific information bleeding across folds) leading to an overestimation of performance. This consideration is particularly important because our features are largely defined by query-biomarker and query-target relationships. To ensure the model demonstrates generalisation rather than memorisation, we treat the random-split result as an optimistic upper bound and introduce an alternative benchmarking scheme tailored to chemogenetic CRISPR data.

**Fig 5.**
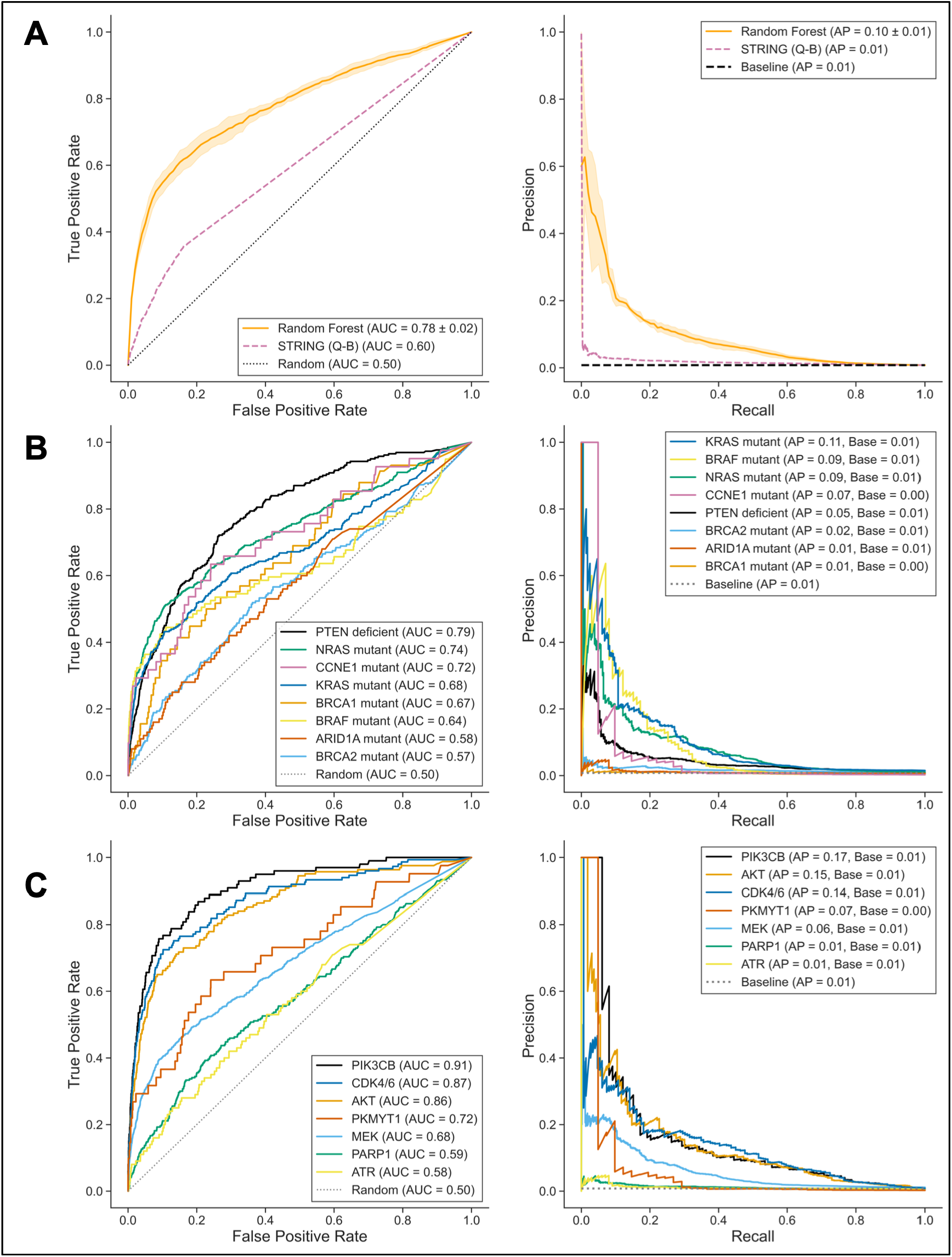
Performance of SL resistance classifiers across distinct evaluation strategies. **A)** Receiver Operating Characteristic (ROC) and Precision-Recall (PR) curves comparing random forest classifier predictions (orange) against single-feature baselines (STRING query-biomarker interaction score, pink and random baseline, grey) in random train-test splits with 5-fold stratified cross-validation. Mean AUROC and average precision (AP) scores with standard deviations across folds are reported in the legend. **B)** ROC and PR curves for the leave-one-biomarker-out evaluation scheme, where models were trained on all but one biomarker group and tested exclusively on the held-out group. **C)** ROC and PR curves for the leave-one-target-out evaluation scheme, where all samples from a given drug target were held out for testing and the classifier trained on the remaining data.

To obtain a more realistic estimate of generalisation and to reduce these potential data-leakage effects, we next evaluated model performance under structured ‘leave-one-group-out’ schemes. To do this, we created three schemes in which all triplets belonging to a given biomarker, target, or SL pair were excluded from training and used exclusively for testing. In the leave-one-biomarker-out evaluation all screens associated with a single biomarker (e.g. PTEN) were used as the test set and all other screens were used for training. In this analysis the best AUROC scores were achieved for PTEN deficiency (Dunn et al., 2022) (AUROC ∼0.79, AP ∼5%) and *NRAS* mutation (Hayes et al., 2019) (AUROC ∼0.74, AP ∼9%) (Fig. 5B). Although performance varied across screens associated with distinct biomarkers, 6/8 biomarkers had an AUROC of 0.60 or greater.

In the leave-one-target-out evaluation (Fig. 5C), all screens associated with a given target (e.g. MEK) were kept as the test dataset and the screens associated with remaining targets used as training data. Performance was strongest for the PIK3CB (AUROC ∼0.91, AP ∼17%), CDK4/6 (AUROC ∼0.87, AP ∼14%) and AKT (AUROC ∼0.86, AP ∼15%) inhibitor screens, and remained above random for PKMYT1 (AUROC ∼0.72, AP ∼7%) and MEK (AUROC ∼0.68, AP ∼6%), with weaker discrimination for PARP1 (AUROC ∼0.59, AP ∼1%) and ATR (AUROC ∼0.58, AP ∼1%). This indicates the model generalises across distinct synthetic lethal contexts, including oncogene-driven vulnerabilities such as CCNE1-PKMYT1. Similarly, in the leave-one-SL-pair-out evaluation (Fig. S6), performance spanned AUROC 0.57-0.91 across the ten pairs, with a mean of 0.72 ± 0.12 and a mean average precision of 9% ± 6%. The strongest were PTEN-PIK3CB (AUROC 0.91, AP 17%), NRAS-CDK4/6 (AUROC 0.87, AP 14%) and PTEN-AKT (AUROC 0.86, AP 15%); the weakest — BRCA2-PARP1 and ARID1A-ATR — still exceeded random expectation (AUROC 0.57-0.58).

Overall, although performance varied across settings, our classifier demonstrated predictive power, achieving AUROC scores above 0.6 in most evaluation settings. These results indicate that, even under stringent leave-one-group-out schemes, the model captures generalisable patterns of resistance rather than biomarker, target or pair specific relationships. Together, these findings suggest that integrating protein-protein interaction, expression, and essentiality features provides a robust and biologically grounded framework for anticipating resistance across diverse synthetic lethal contexts.

### Predicting resistance mechanisms to SL therapies in clinical trials

To assess the translational relevance of our model, we applied our random forest classifier to predict mechanisms of resistance to 10 synthetic lethal therapies currently under active clinical evaluation. These synthetic lethal candidates (biomarker-target pairs) were obtained from a recent review of clinical trials investigating synthetic lethal strategies (Gonçalves et al., 2025) and include inhibitors targeting proteins involved in DNA damage response and genome maintenance, cell-cycle regulation, epigenetic regulation, and receptor tyrosine kinase signalling (Table-2). For each of these SL pairs, we generated predictions of resistance across all protein coding genes in the HGNC reference database, applying the same leave-one-out scheme used above — for each pair, any training screen sharing its biomarker or target was excluded before training, to avoid leakage. This analysis extends beyond published resistance screens and provides a systematic, genome-wide prioritisation of candidate resistance genes for 10 clinically relevant SL therapies (Supplementary File-2). We built an interactive web resource (https://cancergenetics.github.io/sl-resistance-database) that exposes the full prediction output across all pairs. The database supports two complementary query modes: given a biomarker-target pair, users obtain a ranked list of candidate resistance genes across the protein-coding genome; given a gene of interest, users retrieve all SL contexts in which that gene is predicted as a high-confidence resistance modifier. Each entry is annotated with the corresponding clinical trial identifiers and therapeutic agents. The resource is openly accessible and is designed to support experimental prioritisation, biomarker interpretation, and hypothesis generation for synthetic lethal therapies currently under clinical investigation.

**Table-2:** List of synthetic lethal biomarker–target pairs evaluated in clinical trials and used for genome-wide resistance prediction. DDR, DNA damage response; HRD, homologous recombination deficiency; MSI-H/dMMR, microsatellite instability high or mismatch-repair deficiency.

| Chosen biomarker gene | Drug target | Clinical trial ID | Notes on clinical biomarker |
| --- | --- | --- | --- |
| *ATM* | ATR | NCT04657068, NCT04497116, NCT02278250 | DDR gene mutations |
| *BRCA1* | ATR | NCT04905914 | DDR gene mutations |
| *BRCA1* | PARG | NCT05787587 | HRD |
| *BRCA1* | POLQ | NCT05687110, NCT04991480, NCT05898399, NCT06077877, NCT06545942 | HRD |
| *BRCA1* | USP1 | NCT06065059, NCT05240898 | HRD |
| *MSH6* | WRN | NCT05838768, NCT06004245 | MSI-H/dMMR |
| *MTAP* | PRMT5 | NCT05245500, NCT05094336, NCT05732831, NCT06130553 |  |
| *FBXW7* | PKMYT1 | NCT04855656 |  |
| *KRAS* | PLK1 | NCT03829410 |  |
| *CDH1* | ROS1 | NCT03620643, NCT04551495 |  |

Among the SL pairs we made predictions for is MTAP-PRMT5 – a promising SL pair currently undergoing multiple clinical trials. The MTAP-PRMT5 synthetic lethal relationship arises because loss of MTAP leads to accumulation of methylthioadenosine, which inhibits PRMT5 activity, creating a collateral vulnerability that can be therapeutically exploited (Kryukov et al., 2016; Marjon et al., 2016; Mavrakis et al., 2016). Several PRMT5 inhibitors targeting this dependency are currently in early-phase clinical trials for MTAP-deleted tumours (Engstrom et al., 2023; Rodon et al., 2024). A loss-of-function CRISPR screen to dissect the MTAP-PRMT5 synthetic lethality, identifying genes whose loss conferred both increased sensitivity and resistance to PRMT5 inhibitors in an MTAP null setting, was recently published (Knoll et al., 2025). This screen used a focused library targeting the ‘druggable’ genome (4,547 genes) rather than a genome-wide screen and so was not included in our initial training. The top resistance hit identified was CARM1, which encodes PRMT4 and has been reported to physically interact with PRMT5, consistent with our hypothesis that resistance is often driven by functionally related genes (Nie et al., 2018; Yuan et al., 2022). To assess the overall utility of our predictions in this setting, we followed the same workflow as the other drug resistance screens and identified the top 100 resistance causing genes from this screen as the resistance labels. Our classifier demonstrated good performance in separating resistance from non-resistance genes with an AUROC score of ∼0.71 and an AP of ∼7% (Fig. 6A, Fig. S7A). The model outperformed STRING protein-protein interaction scores between query and biomarker (AUROC of ∼0.57, AP of ∼3%). Importantly, the ability of the classifier to recover meaningful resistance signals in a biologically distinct SL context suggests that the model captures generalisable principles of drug resistance rather than pair-specific artefacts.

**Fig 6.**
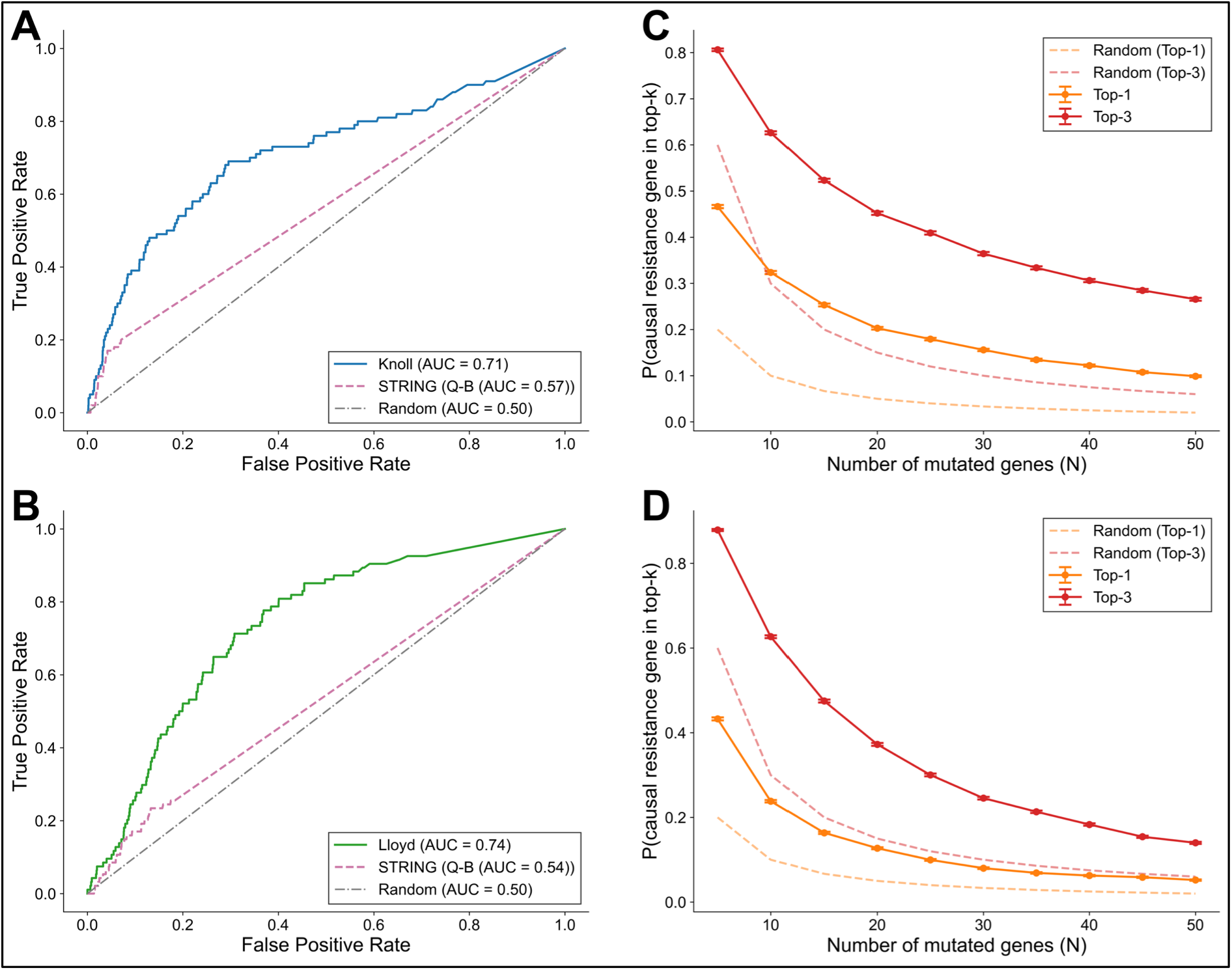
Evaluation of the prediction of resistance to PRMT5 inhibitors and ATR inhibitors. **A**) ROC curve evaluating predictions for resistance to PRMT5 inhibitors in MTAP-deleted cancer cells (Knoll et al., 2025). Our classifier is shown in blue and compared against STRING interaction scores (pink dashed) and random expectation (grey dash-dotted). B) ROC curve for the ATM-deficient ATRi screen. The random forest classifier (green) was trained on the main dataset excluding ARID1A-ATR pair. Baselines as in A. C) Empirical top-k analysis of the model’s ability to prioritise causal resistance genes among N mutated candidates, for the MTAP-PRMT5 and D) ATM-ATR screens. For each value of N (x-axis), we estimate the probability that the causal resistance gene appears in the top-1 or top-3 ranked predictions (solid lines, y-axis). Model performance is compared against a random selection baseline (dashed lines). Error bars represent standard error across 20,000 simulation trials.

Another SL pair undergoing clinical trials for which we made predictions is ATM-ATR. Loss of ATM, a key kinase in the DNA damage response, compromises the cellular response to double-strand breaks and collapsed replication forks, increasing dependency on ATR-mediated replication stress resolution (Menezes et al., 2015; Reaper et al., 2011; Schmitt et al., 2017; Toledo et al., 2011), establishing a synthetic lethal vulnerability exploitable with ATR inhibitors. ATR inhibitors, including gartisertib (M4344), are currently being evaluated in multiple clinical trials in ATM-deficient tumours (Burris et al., 2024). A genome-wide CRISPR knockout screen was previously published to identify genes whose loss confers resistance to ceralasertib (AZD6738), an ATR inhibitor that is itself being evaluated in clinical trials in ATM-altered tumours (NCT04564027, NCT03682289), in an ATM-deficient background in mouse embryonic stem cells (Lloyd et al., 2021). Because this study was performed in mouse cells, it was not included in the datasets used to train our classifier. Nonetheless, we reasoned that we may be able to make predictions for resistance by mapping our predictions for human genes to their mouse orthologs. Because a screen with ATR inhibitor (as ARID1A-ATR) (Llorca-Cardenosa et al., 2022) was present in our training data, the ATM-ATR predictions in our database were generated with the ARID1A-ATR screen excluded from training, following the leakage-exclusion scheme described previously. We used these database predictions directly as above and compared them with the resistance labels of the screen. Our model achieved an AUROC of ∼0.74 and AP of ∼2% (Fig. 6B, Fig. S7B) in this screen and it outperformed STRING protein-protein interaction scores between query and biomarker (AUROC ∼0.54, AP ∼1%). These results further support the generalisability of our framework across distinct SL contexts, drug classes and species.

### Prioritising resistance genes among mutated genes

The analyses presented so far demonstrate how well our classifier performs when identifying resistance genes from among the full set of ∼19,000 protein coding genes in the genome, an extremely challenging scenario given the rarity of resistance genes. However, as additional SL therapies enter the clinic, other use cases are likely to be of increasing importance. In particular, when patients develop resistance to an SL therapy and the genomes of resistant samples are sequenced our predictions may be of use in prioritising the genetic variants most likely to cause resistance. This scenario is analogous to prioritisation of causal variants from exome or genome sequencing in rare Mendelian disease, where tools such as Exomiser rank candidate genes from a patient’s variant list using phenotypic associations and protein-interaction-network evidence (Smedley et al., 2015). Here, the goal is not to identify a phenotype-causing gene from the entire genome but rather from the smaller set of genes mutated in a specific patient. To model this, we ranked one “causal” resistance gene against a background of N−1 non-resistance genes from both the MTAP-PRMT5 and the ATM-ATR screens and measured how often it placed within the top-1 or top-3 (Monte Carlo simulation; see Methods). Our model consistently outperformed a random classifier baseline across all values of N in both screens (Fig. 6C, 6D). As expected, the ranking probability decreases with increasing N, reflecting the increasing difficulty of prioritisation as the candidate space grows. These results suggest that the model captures meaningful signals for resistance gene prioritisation and can robustly elevate causal variants above a pool of background mutations. The empirical top-k framework also mimics a realistic interpretation setting, in which researchers seek to analyse candidate mutations from a limited list of alterations in individual tumour samples.

Taken together, these results demonstrate that the integrative feature set and model architecture generalise beyond the original training dataset to accurately capture resistance determinants in biologically distinct SL settings.

## Discussion

As more synthetic lethal therapies enter the clinic, it will be necessary to understand the mechanisms underlying therapeutic resistance. Here, we introduce the first machine learning framework to systematically predict resistance across diverse SL pairs. Our approach builds on large-scale CRISPR-Cas9 chemogenetic screens and is, to our knowledge, the first to propose a benchmarking scheme specifically designed to evaluate generalisation of predictive models of SL resistance.

Analyses of published chemogenetic screens revealed that genes physically or functionally connected to SL biomarkers and targets are significantly enriched among resistance genes. This enrichment, observed across nearly all screens, suggests that resistance is not random but follows relationships defined by molecular interaction networks. These findings extend previous observations that functionally linked genes are more likely to display synthetic lethality (Benstead-Hume et al., 2019; Costanzo et al., 2016; De Kegel et al., 2021; Kelley & Ideker, 2005; Lord et al., 2020) and highlight network proximity as key to therapeutic escape. Building on these observations, we reasoned that integrating PPI with other molecular features could yield a more generalisable model of resistance. PPI-based features highlight the functional proximity of query genes to SL biomarkers and targets, essentiality-derived features capture context-dependent dependencies across cell lines, and expression features reflect transcriptional co-regulation. When combined within an integrative random forest model, these features consistently outperform any individual feature.

Although our classifier consistently showed predictive power across biomarker, target, and SL-pair benchmarking schemes, performance varied between contexts. These differences likely reflect both biological and experimental sources of variability. Screens performed in different cell lines often yield only partially overlapping resistance hits (Fig. S1), underscoring that SL resistance mechanisms can be highly context-specific. Moreover, differences in screen design, such as library composition, drug exposure time, and computational processing, may contribute to variable signal-to-noise ratios and affect apparent model performance. For instance, the lower AUROCs observed in certain biomarkers may reflect differences in screen sensitivity rather than an intrinsic lack of predictability (Fig. 5B).

Our models retained predictive power across leave-one-biomarker, leave-one-target, and leave-one-SL-pair-out schemes, in which all data from a given biomarker, target, or SL pair were held out to reduce data leakage and test extrapolation to unseen contexts. The successful application to independent MTAP-PRMT5 and ATM-ATR case studies demonstrates generalisation beyond the original training datasets. These results indicate that the model captures fundamental principles of resistance rather than pair-specific artefacts. Nevertheless, precision values remain low, reflecting both the extreme class imbalance (resistance genes are <1% of screened genes) and the heterogeneity of resistance mechanisms. A further current limitation is that resistance labels are aggregated across the heterogeneous screens described above, which may introduce label noise into the training set. Future work incorporating background-specific models that account for cell-line or lineage-dependent features may help separate biological heterogeneity from technical variation and further improve predictive accuracy. In addition, the number of currently profiled SL pairs remains modest. As more pairs are subject to chemogenetic resistance screens we anticipate the predictive accuracy of our model will increase. Perhaps more important will be the availability of resistance mechanisms identified in patients rather than cell lines. Currently this data is only available for PARP inhibitor resistance, but as more SL therapies enter the clinic this is likely to change. While further work is needed to expand feature types, improve precision, and validate predictions experimentally, this study provides a foundation for predictive approaches in SL resistance.

## Materials and Methods

### Data Collection and Labelling

We curated genome-wide CRISPR-Cas9 chemogenetic resistance screens across nine publications and spanning ten unique SL combinations (Table 1) from the published literature that evaluated resistance to targeted therapies in synthetic lethal (SL) contexts. Where a study screened several cell lines, inhibitors of the same target, or clonal derivatives of one line, these were aggregated into a single gene-level resistance dataset per study and pair, giving thirteen screen datasets. Screens were included if they reported gene-level resistance ranking in the presence of a drug, with a defined biomarker-target SL pair, and were performed in human cell lines. Where an inhibitor targets multiple paralogs, the target was represented by all targeted genes (i.e. pan-AKT inhibitor: AKT1/AKT2/AKT3; MEK inhibitors: MAP2K1/MAP2K2; CDK4/6 inhibitor: CDK4/CDK6). Screen results were obtained from the supplementary materials of the original manuscripts, and the resulting gene lists were cross-checked against the CRISPR library reported in each study to account for potential missing entries. For each screen, genes identified as resistance hits were labelled as “resistance”, while all other genes were labelled as “non-resistance”. If authors reported a clear statistical threshold for defining resistance hits, we adopted this threshold directly; otherwise, we labelled the top 100 genes ranked by the study’s primary scoring metric. When multiple studies screened the same SL pair (e.g., NRAS-MEK1/2, BRCA1-PARP1, KRAS-MEK1/2), results were aggregated. For aggregated screens, a union rule was applied such that a gene was labelled “resistance” if it was classified as resistance in at least one included study.

All query gene symbols were resolved against the HGNC complete set (Seal et al., 2023) (downloaded on April 12, 2024) using a fixed precedence: each symbol was matched first to an approved HGNC symbol; if no approved match existed, to a previous symbol; and finally to an alias symbol. The first matching approved gene was retained, and symbols matching none were left unmapped. This ordering ensures that a symbol serving as the current approved name for one gene is always assigned to that gene, and that a former official (previous) symbol takes precedence over an informal alias. Each resolved gene was then assigned its corresponding HGNC ID, Entrez Gene ID and Ensembl gene ID (ENSG).

### Protein-Protein Interaction Overlap

To assess whether resistance genes were enriched among protein-protein interaction (PPI) partners of synthetic lethal pairs, we compiled PPI data from STRING (Szklarczyk et al., 2023) (downloaded on December 7, 2023, version 12.0) and BioGRID Physical (Oughtred et al., 2021) (downloaded on November 28, 2024, version 4.4.241). For each biomarker and target gene in our dataset, we retrieved their reported interaction partners from each database. We then tested the overlap between these interactors and the resistance genes identified in each CRISPR screen. Statistical enrichment was calculated using Fisher’s exact test, reporting odds ratios and p-values. Analyses were performed separately for STRING (medium-confidence network, score ≥ 0.4) and BioGRID physical interactions and were repeated across individual screens as well as in aggregated datasets.

To determine whether the observed overlap between resistance genes and the PPI partners of SL biomarkers and targets exceeded that expected by chance, we also performed a degree-preserving network randomisation analysis using the configuration model (Britton et al., 2006; Molloy & Reed, 1995). Across all SL pairs, we counted the resistance genes connected to either the biomarker or target in the real PPI network. Self-loops were removed from the real network prior to degree extraction to ensure an accurate degree sequence. We then generated 1,000 degree-matched random networks using the Python package NetworkX (version 3.2.1) (Hagberg et al., 2008), retaining the exact node degree distribution of the STRING (medium-confidence) and BioGRID physical interaction networks while randomising edge assignments. Parallel edges and self-loops introduced during randomisation were removed to maintain simple graph structure. For each randomised network, we counted the total number of resistance genes (pooled across all SL pairs) connected to their biomarker or target, producing an empirical null distribution under a network with identical degree structure but random wiring. The overlap observed in the real STRING and BioGRID networks was then compared to the distribution of random overlaps, and the 10th-highest and the 50th-highest random values were used as reference thresholds for p < 0.01 and p < 0.05, respectively.

### Preprocessing and Feature Engineering

All resistance screens were imported into a main dataset by parsing individual screen outputs and appending study-specific metadata, including screen identifier and SL pair labels. Synthetic lethal pairs were decomposed into their constituent biomarker and target genes. For multi-paralogue targets (defined above), each targeted gene was stored as Target1, Target2 and Target3 as applicable (Target2/Target3 left undefined for two– or single-gene targets). Each target-based feature was computed separately against each defined target and combined into a single query-target feature by retaining the largest-absolute-magnitude value, or the available value where one was missing. Column names were harmonised (e.g., “Query” for candidate resistance genes, “Class” for resistance labels), and the dataset was reordered to place identifiers and annotations in a consistent format for downstream feature engineering and model training. The features are predominantly quantitative and can be grouped into three broad categories, expression-, essentiality– and protein-protein interaction based (Supplementary File-1). To account for the functional nature of biomarker alterations, we annotated each biomarker gene using the Cancer Gene Census (CGC) (Sondka et al., 2018). Mutations were classified as either loss-of-function (typical of tumour suppressor genes) or activating (typical of oncogenes). A categorical feature was then defined where loss-of-function mutations were encoded as 1 and activating mutations as 0.

#### Protein-protein interaction-based features

To quantify network-level proximity between query genes and synthetic lethal partners, we derived four “shared protein-protein interactors” features from the STRING (medium-confidence, score ≥ 0.4) and BioGRID physical interaction networks described above. For each query-biomarker and query-target pair, we compared their sets of interactors and assessed the significance of their overlap using Fisher’s exact test; the resulting signed –log10 p values served as the feature scores.

We also derived direct PPI features from STRING and BioGRID. From STRING, the combined interaction score was retrieved for each query-biomarker and query-target pair. These scores provide a continuous measure of interaction confidence derived from multiple evidence channels. For BioGRID, we assigned a binary feature indicating whether a direct physical interaction between the query gene and the biomarker or target gene was reported (1 = interaction, 0 = no interaction). Together, these features capture direct molecular associations between candidate resistance genes and SL partners across complementary interaction databases.

#### Expression-based features

Gene expression features were derived from Genotype-Tissue Expression (GTEx) bulk tissue expression data (GTEx Consortium, 2020) (downloaded on February 5, 2025, version 10, RNASeQCv2.4.2). Expression profiles for query, biomarker, and target genes were extracted and filtered to exclude missing values. For each query-biomarker and query-target pair, we calculated the Spearman correlation coefficient of their expression levels across GTEx samples, yielding two co-expression features. In addition, we computed the average expression level and the variance of expression for each query gene across GTEx samples.

#### Essentiality-based features

Essentiality features were derived from genome-scale CRISPR dependency profiles reported by the Cancer Dependency Map (DepMap) (Behan et al., 2019; Meyers et al., 2017) 23Q4 dataset (DepMap, 2023). For each query-biomarker and query-target pair, we calculated the Pearson correlation between their essentiality profiles across DepMap cell lines. For each query gene individually, we also computed the average essentiality score and the variance of essentiality scores across cell lines. In addition, we calculated the percentage of cell lines in which a query gene is classified as essential, using a predefined threshold of gene effect score < –0.6.

### Model Optimisation and Hyperparameter Selection

To identify the optimal parameters for our random forest classifier, we performed an extensive grid search using the GridSearchCV function from scikit-learn, coupled with a leave-one-group-out (LOGO) cross-validation scheme where groups corresponded to SL pairs. Model performance was evaluated using both average precision (AP) and AUROC, with AP used as the primary metric for selecting the best hyperparameter configuration, given the strong class imbalance between resistance and non-resistance genes.

We explored a hyperparameter space spanning n_estimators (100, 300, 500, 1,000), max_features (“sqrt”, 0.3, 0.5, and None), max_depth (None, 10, 20, 30), min_samples_leaf (1, 4, 8), and class_weight (None, “balanced”, {0: 1, 1: 5}, and {0: 1, 1: 10}). The search was conducted across all ten SL pairs. The final configuration was selected by ranking models by their mean AP and AUROC across all LOGO folds and averaging the two ranks, so the chosen model performed well on both metrics. Using this criterion, the optimal settings were identified as n_estimators = 300, max_features = “sqrt”, max_depth = None (unlimited), min_samples_leaf = 4, and class_weight = {0: 1, 1: 10}.

### Model Training and Design

We trained a random forest classifier (scikit-learn) to predict resistance labels from a total of 18 features in the main dataset, using the optimal hyperparameters described above. For the random-split evaluation, we used 5-fold stratified, shuffled cross-validation on the full dataset (each fold an 80:20 train/test split) to preserve class balance within folds. We computed AUROC and AP per fold; mean curves were obtained by interpolating TPR over a common FPR grid (ROC) and precision over a common recall grid (PR), followed by averaging across folds. We report the mean ± standard deviation of AUROC and AP across folds. The baseline precision for PR plots was defined as the positive class prevalence in the corresponding test data. To benchmark against single-feature baselines, we evaluated the STRING interaction score between query and biomarker as a standalone scorer by computing the ROC and PR curves and associated AUROC/AP on the same labels (no model fitting).

To assess generalisation while minimising data leakage, we used three hierarchical leave-out schemes. In each, all instances sharing a given target (leave-one-target-out), biomarker group (leave-one-biomarker-out; eight groups: PTEN, BRCA1, BRCA2, CCNE1, NRAS, KRAS, BRAF, ARID1A), or SL pair (leave-one-SL-pair-out) were held out for testing while the model was trained on the remainder with the same hyperparameters; AUROC and AP were computed on each held-out group.

### Clinical trials model application and prediction database

To extend our resistance predictions beyond the training screens, we identified clinically relevant synthetic lethal biomarker-target pairs from a recent systematic review of SL-directed clinical trials (Gonçalves et al., 2025). From this resource, we curated 10 unique biomarker-target pairs currently under active clinical investigation (Supplementary File-2), spanning 22 clinical trials across Phase I and Phase II settings. For each of the 10 biomarker-target pairs, the query set comprised all protein-coding genes in the HGNC reference database, each already annotated with its HGNC, Entrez and Ensembl identifiers. The same 18 molecular features were then computed for every query gene relative to the corresponding biomarker and target, using the feature-engineering pipeline applied to the training data. The trained random forest classifier was then applied to each dataset to generate genome-wide resistance scores (Supplementary File-2), which were subsequently used to populate the prediction database described below.

To provide open access to the genome-wide resistance predictions for the 10 clinical-stage synthetic lethal pairs, we built an interactive web application (https://cancergenetics.github.io/sl-resistance-database/). The resource was implemented as a client-side static HTML/CSS/JavaScript application with no server-side computation, ensuring unrestricted accessibility without infrastructure dependencies. All prediction data were pre-processed offline using a custom Python pipeline (available at the associated GitHub repository). These genome-wide predictions (approximately 19,000 genes per pair; 191,290 scored gene-SL pair entries in total) were ranked in descending order of predicted resistance score, with higher scores indicating a stronger predicted likelihood of conferring resistance. The processed predictions were serialised into a set of pre-built JSON files, partitioned into therapy-pair-indexed files for pair-centric queries and MD5-sharded gene bucket files for gene-centric queries, enabling real-time search and filtering without server-side computation. The interface supports two complementary query modes: users may select a biomarker-target pair to retrieve the complete ranked gene list for that SL context, or search by gene symbol to identify all 10 SL pairs in which that gene is predicted as a resistance candidate and its corresponding rank. Clinical trial metadata, including trial identifiers, phase, and therapeutic agent, were compiled from the review of SL clinical trials (Gonçalves et al., 2025) and linked to each SL pair within the resource.

### Model Validation and Case Study

To evaluate the generalisability of our framework, we applied the random forest classifier to an independent pooled single-gene CRISPR knockout screen assessing fitness under PRMT5 inhibition in MTAP-deleted cancer cell lines (LU99 and SW1573) (Knoll et al., 2025). Gene-level log fold change (logFC) values were retrieved from the supplementary materials, and an average differential log fold change score (AvgDiff) was computed as the mean of the drug-versus-DMSO differential log fold change across the two cell lines. Paralog-targeting (digenic) guides and control sgRNAs were excluded prior to labelling. Genes with positive AvgDiff were ranked in descending order, and the top 100 ranked genes were labelled as resistance, with all others considered non-resistance (n = 4,547 genes total). Because MTAP-PRMT5 is one of the ten clinically evaluated biomarker-target pairs scored above, we took the genome-wide MTAP-PRMT5 resistance predictions directly and matched them to the screen genes by HGNC identifier, rather than re-deriving features or retraining the model. Genes absent from that prediction — pseudogenes outside the protein-coding query set, and protein-coding genes lacking computable features (e.g., no GTEx expression data), all of which were non-resistance — were excluded, leaving 4,528 genes for evaluation. The predicted resistance scores were then compared against the experimental resistance labels to assess concordance.

For further validation, we applied the random forest classifier to a genome-wide CRISPR knockout resistance screen using AZD6738 (ceralasertib, an ATR inhibitor) in ATM-deficient mouse cells (Lloyd et al., 2021). Resistance genes were taken from the genome-wide positive-selection analysis of the high-dose (IC90) ceralasertib screen, the primary screen in the original publication. Genes were ranked by positive-selection rank as in the original study, and the top 100 were labelled as resistance, with all others considered non-resistance. As the screen was conducted in mouse cells, gene symbols were mapped to human orthologs using the Ensembl BioMart ortholog table (Dyer et al., 2025; Smedley et al., 2009) (release 114, April 2025) as the primary source, with exact-match, previous-symbol and alias lookups against the HGNC complete set (Seal et al., 2023) (downloaded on April 12, 2024) as sequential fallbacks; unmapped genes were excluded. Because ATM-ATR is one of the ten clinically evaluated pairs scored above, we took the genome-wide ATM-ATR resistance predictions directly and matched them to the screen’s human-ortholog genes by HGNC identifier, rather than re-deriving features or retraining the model. After excluding genes with missing features and deduplicating, 15,613 unique query genes were retained for evaluation (94 of the top 100 resistance genes), and the predicted resistance scores were compared against the experimental resistance labels.

### Empirical Top-k Resistance Gene Prioritisation Analysis

Prioritisation performance was assessed by Monte Carlo simulation over 20,000 independent trials. In each trial, one positive gene (known resistance gene) was drawn uniformly at random, and N−1 negative genes were sampled from the non-resistance pool of that screen. The N candidates were ranked by predicted resistance probability, and a trial was counted as a hit if the true resistance gene ranked within the top-k. This procedure was repeated across N ∈ {5, 10, 15, 20, 25, 30, 35, 40, 45, 50} and k ∈ {1, 3}. The empirical hit rate across trials estimated P(causal resistance gene in top-k), with uncertainty quantified as the standard error of the mean across trials. The random classifier baseline for each (N, k) combination is k/N. All simulations used a fixed random seed for reproducibility. This analysis was applied to both validation screens: MTAP-PRMT5 screen (Knoll et al., 2025) and the ATM-ATR screen (Lloyd et al., 2021), using the same predictions and resistance labels described above.

### Software and statistical analysis

All analyses were performed in Python (3.12.9), using pandas (2.1.3) (McKinney, 2010) and NumPy (1.26.4) (Harris et al., 2020) for data handling and SciPy (1.11.4) (Virtanen et al., 2020) for statistical tests (Fisher’s exact test and Mann-Whitney U test). Random forest models and performance metrics (AUROC and AP) were computed with scikit-learn (1.6.1) (Pedregosa et al., 2012), and figures were generated with Matplotlib (3.8.2) (Hunter, 2007); all ROC and precision-recall plots include random and prevalence (positive-class) baselines for reference. Protein-protein interaction networks were visualised in Cytoscape (3.10.2) (Shannon et al., 2003).

## Declarations

### Ethics approval and consent to participate

Not applicable.

Clinical trial number: not applicable.

### Availability of data and material

All analysis code, curated screen tables, and figure-reproduction scripts are available at https://github.com/cancergenetics/SL_resistance. Genome-wide predictions for the ten clinical-stage SL pairs are accessible at https://cancergenetics.github.io/sl-resistance-database/. The study used public data — STRING v12.0, BioGRID v4.4.241, GTEx v10, DepMap 23Q4, HGNC, Cancer Gene Census, and Ensembl BioMart (release 114); CRISPR screens were obtained from the original publications’ supplementary materials (Table 1).

### Consent for publication

Not applicable.

### Authors’ contributions

Conceptualization: CJR, MY

Data curation: MY

Formal analysis: MY

Funding Acquisition: CJR

Methodology: MY

Software: MY

Supervision: CJR

Visualization: MY

Writing – original draft: MY, CJR

Writing – review & editing: MY, CJL, CJR

All authors read and approved the final manuscript.

## Acknowledgements

We thank the members of the Ryan lab for careful reading of the manuscript.

## SUPPLEMENTARY FIGURES

**Fig S1.**
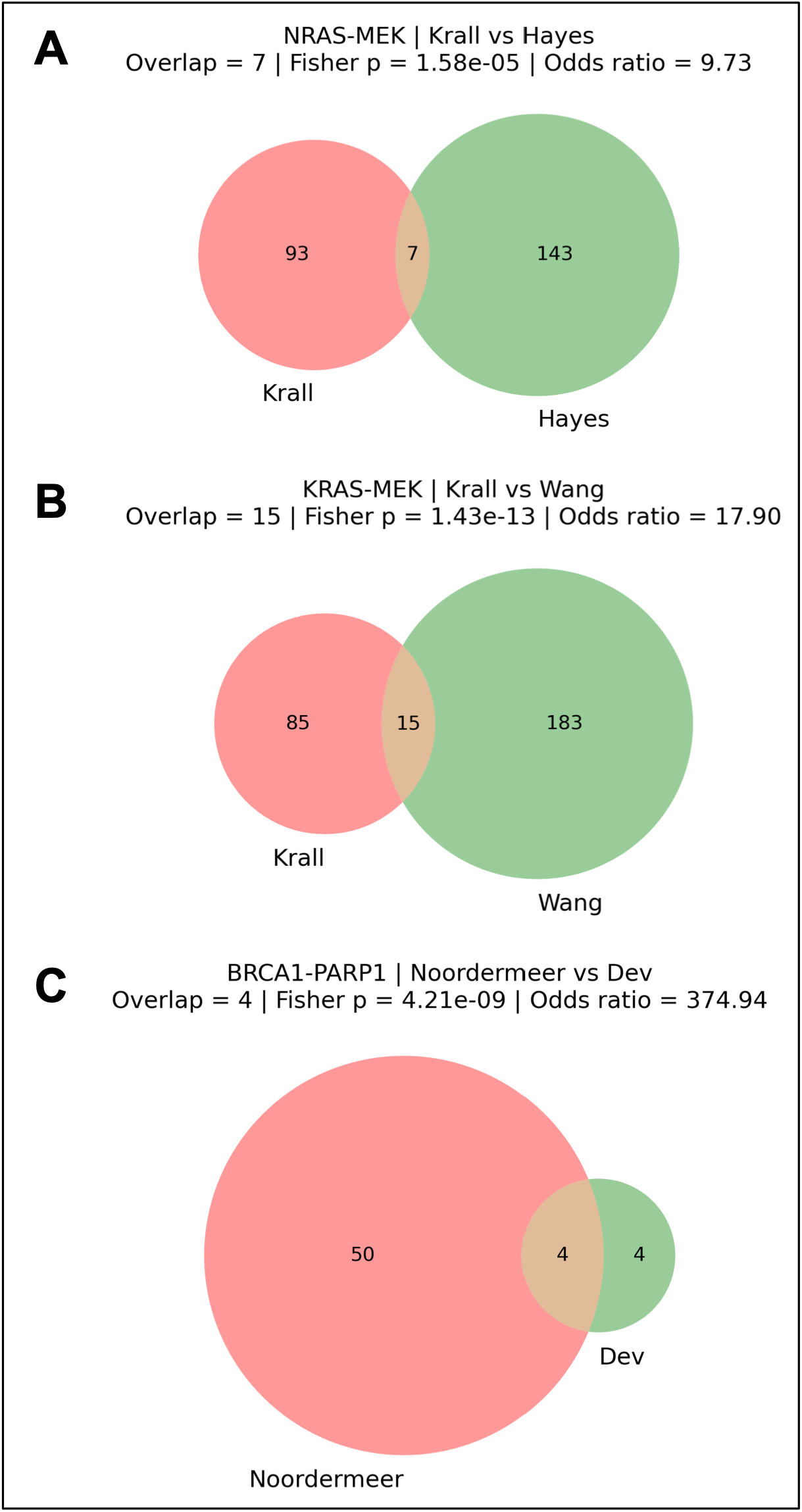
Overlap between hits identified in screens for the same biomarker/target pair. Venn diagrams show the overlap of resistance hits identified in genome-wide CRISPR resistance screens from different studies targeting the same SL pairs. (A) *NRAS-MEK* screens from (Krall et al., 2017) and (Hayes et al., 2019) (B) *KRAS-MEK* screens from (Krall et al., 2017) and (Wang et al., 2017) (C) *BRCA1-PARP1* screens from (Noordermeer et al., 2018) and (Dev et al., 2018). Numbers indicate unique and overlapping resistance hits, with statistical significance of overlap assessed using Fisher’s exact test (p values) and odds ratios.

**Fig S2.**
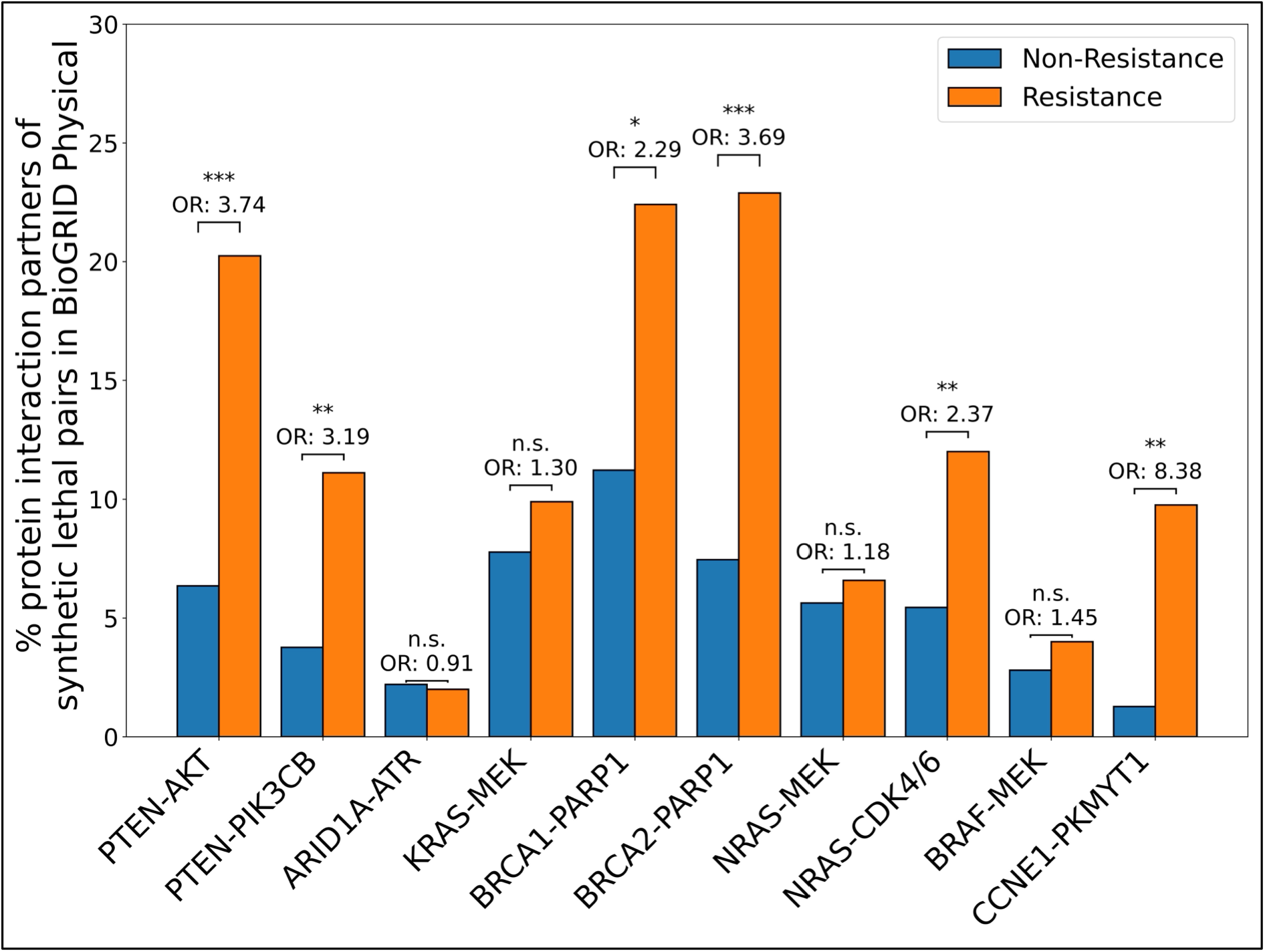
Bar plots of percentages of overlapped protein-protein interaction (PPI) partners in BioGRID physical database among resistance and non-resistance genes. Percentages of resistance (orange) and non-resistance (blue) genes that overlap with known PPI partners of each synthetic lethal (SL) pair in the BioGRID physical interaction database. Statistical enrichments were calculated using Fisher’s exact test, reporting odds ratios and p-values (***p < 0.001, **p < 0.01, *p < 0.05, n.s. = not significant).

**Fig S3.**
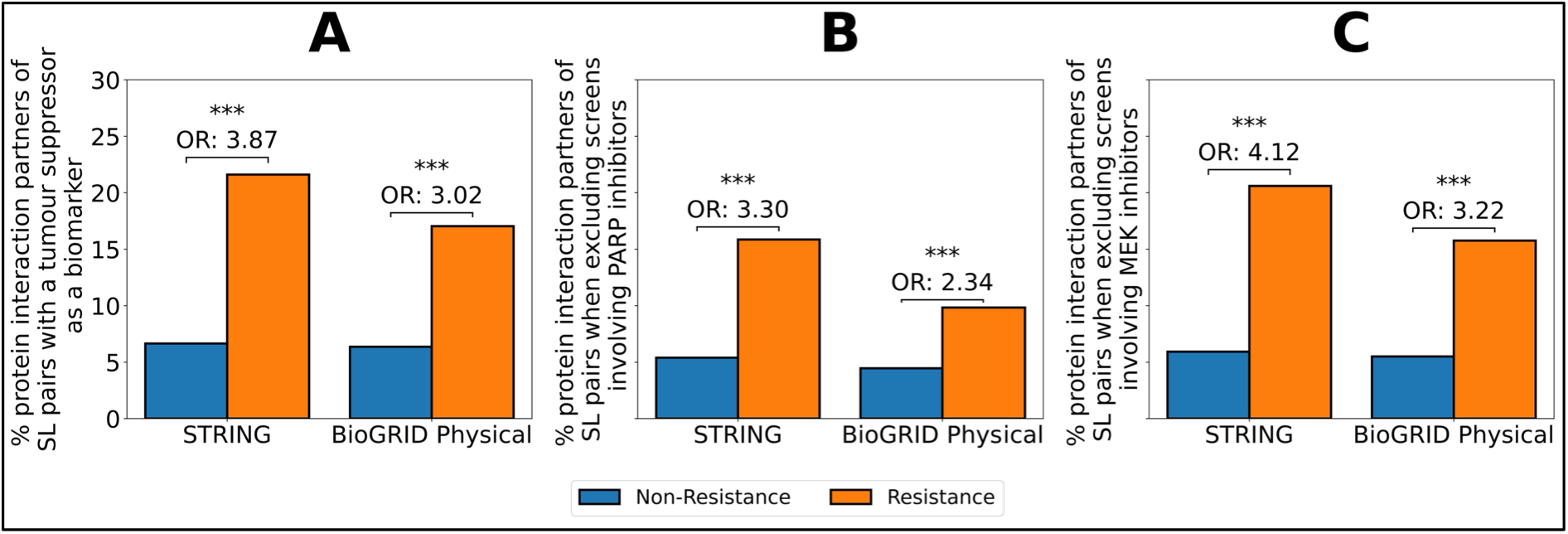
Protein-protein interaction partners among resistance genes across subsets of synthetic lethal (SL) screens. Bar plots show the percentage of resistance (orange) and non-resistance (blue) genes overlapping with the protein interaction partners of SL pairs, as assessed using STRING interaction scores (medium-confidence, ≥ 0.4) and BioGRID physical interactions. (A) SL pairs with a tumour suppressor gene as the biomarker. (B) SL pairs excluding PARP inhibitor screens. (C) SL pairs excluding MEK inhibitor screens. Odds ratios (OR) and statistical significance (Fisher’s exact test, ***p < 0.001) are indicated above each comparison.

**Fig S4.**
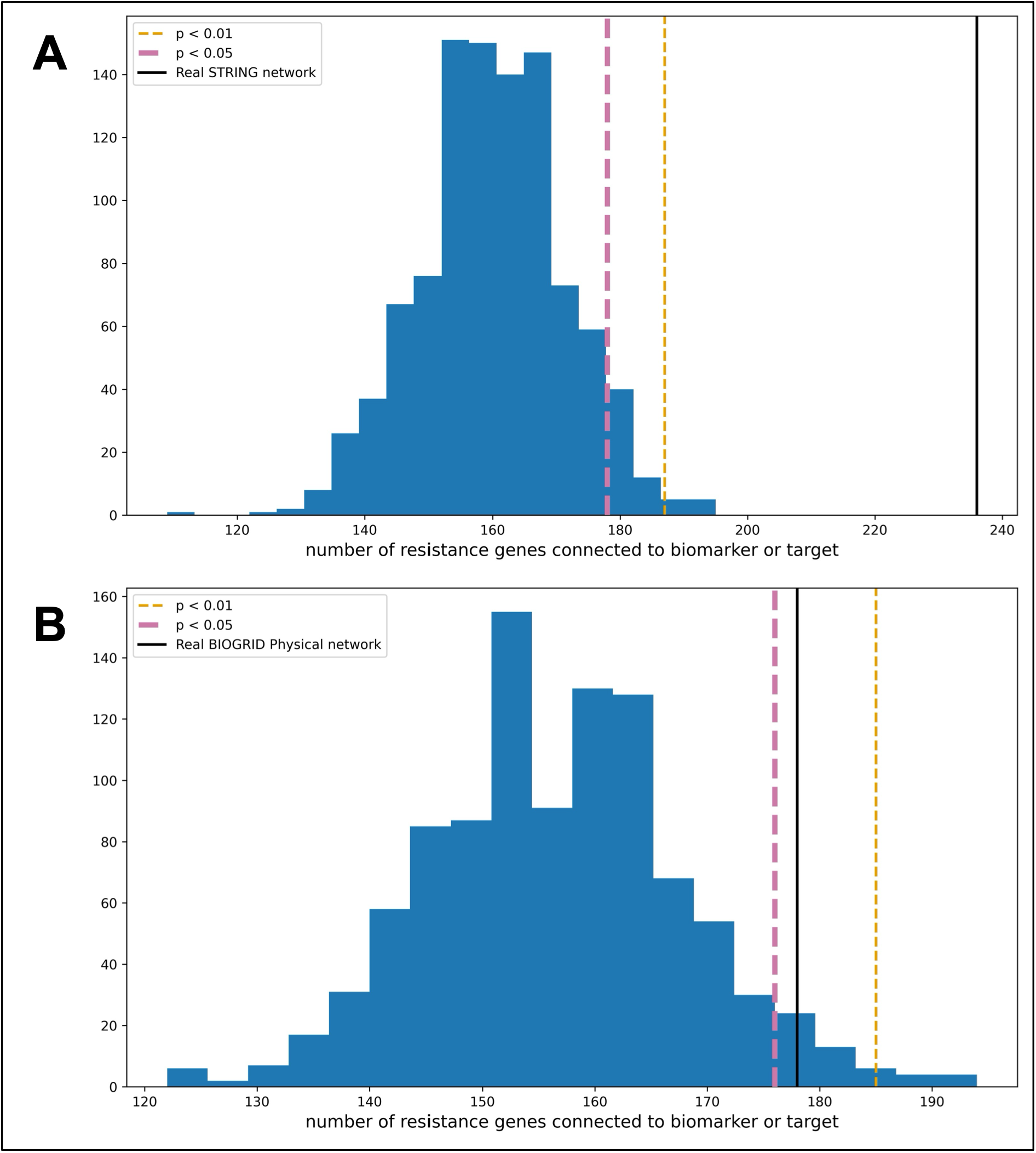
Random network analysis of resistance gene connectivity to synthetic lethal partners. **(A)** Histogram showing the distribution of the number of resistance genes connected to either the biomarker or target across 1,000 degree-preserving randomisations of the STRING medium-confidence PPI network. Vertical lines mark the real STRING overlap (black), the p < 0.01 threshold corresponding to the 10th-highest random value (yellow), and the p < 0.05 threshold corresponding to the 50th-highest random value (pink). **(B)** Equivalent analysis using 1,000 degree-preserving randomisations of the BioGRID physical interaction network. Vertical lines indicate the real BioGRID physical overlap (black), the p < 0.01 threshold (yellow), and the p < 0.05 threshold (pink).

**Fig S5.**
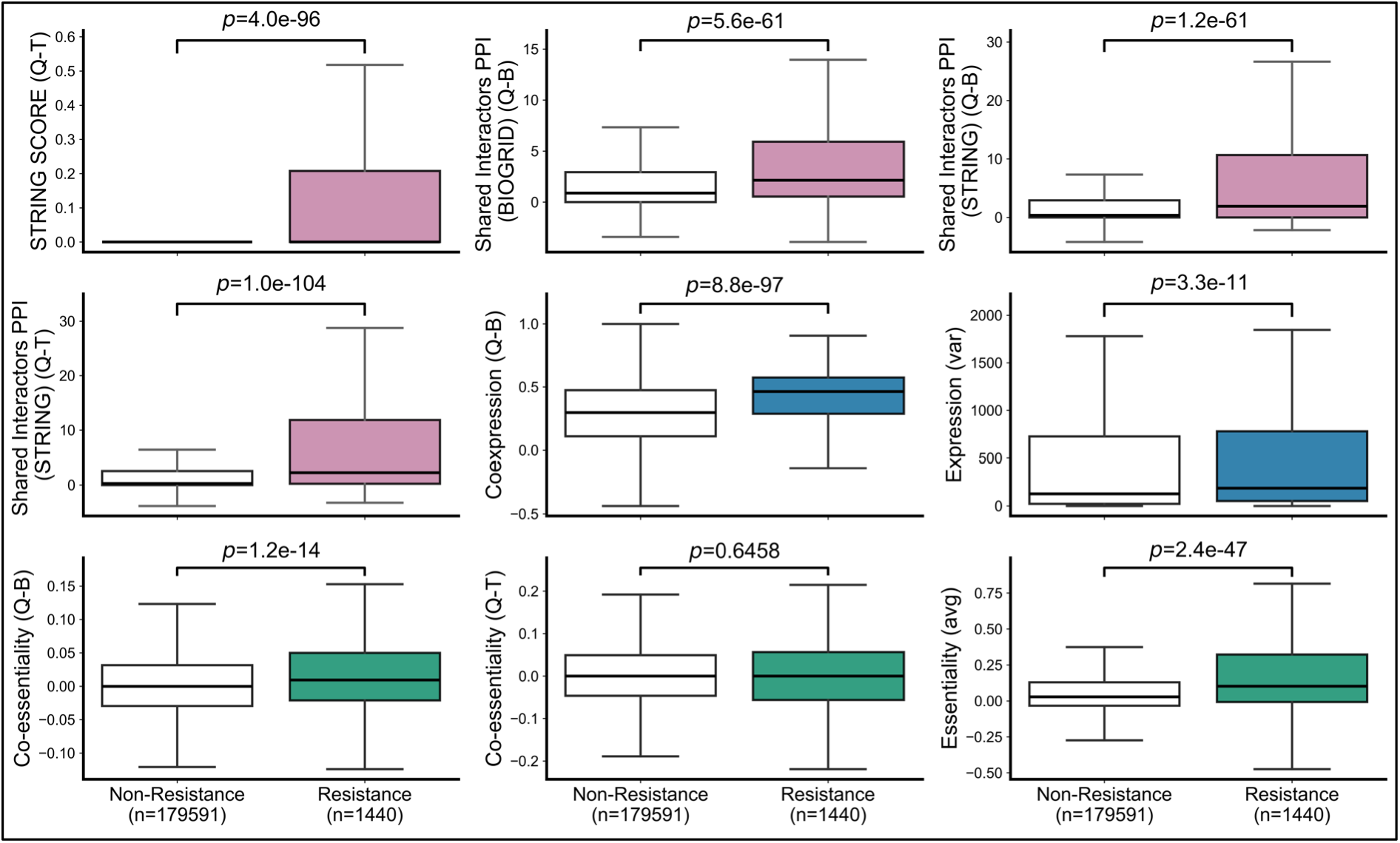
Distribution of additional expression-, essentiality-, and PPI-based features in resistance versus non-resistance genes. Boxplots display the distributions of individual features for resistance (n = 1,440) and non-resistance genes (n = 179,591). Features include STRING protein-protein interaction (PPI) scores, shared interactors from STRING and BioGRID, co-expression correlations, query gene expression variance, co-essentiality correlations, and essentiality averages. Statistical significance between groups was assessed using the Mann-Whitney U test, with p values shown above each comparison.

**Fig S6.**
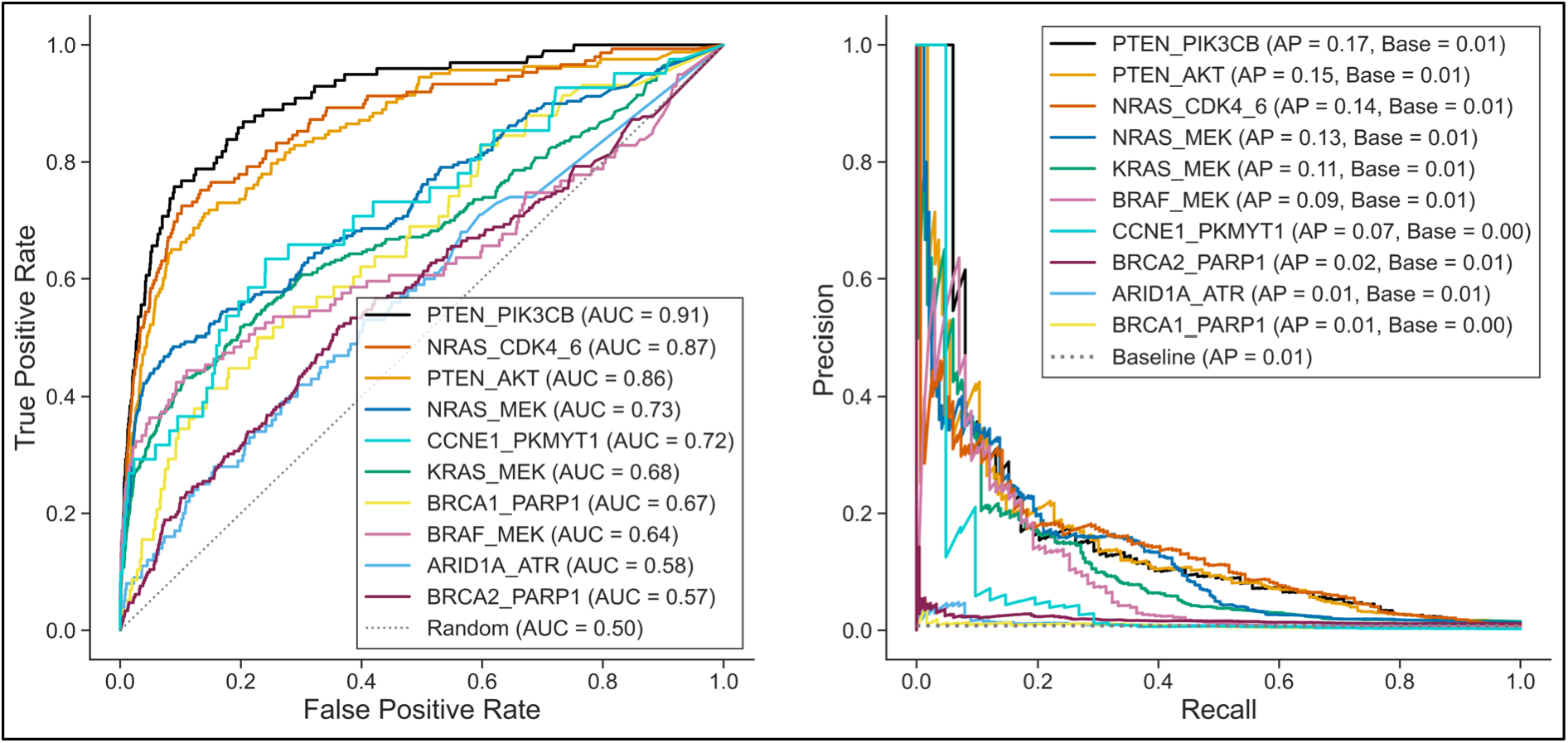
Leave-one-SL-pair-out evaluation of random forest classifier performance. Receiver operating characteristic (ROC, left) and precision-recall (PR, right) curves showing performance of the random forest classifier when individual synthetic lethal (SL) pairs were excluded from training and used exclusively for testing. Each curve corresponds to a distinct SL pair: PTEN-PIK3CB (black), NRAS-CDK4/6 (red), PTEN-AKT (orange), NRAS-MEK (dark blue), BRAF-MEK (pink), ARID1A-ATR (light blue), KRAS-MEK (green), CCNE1-PKMYT1 (cyan), BRCA1-PARP1 (yellow), and BRCA2-PARP1 (purple).

**Fig S7.**
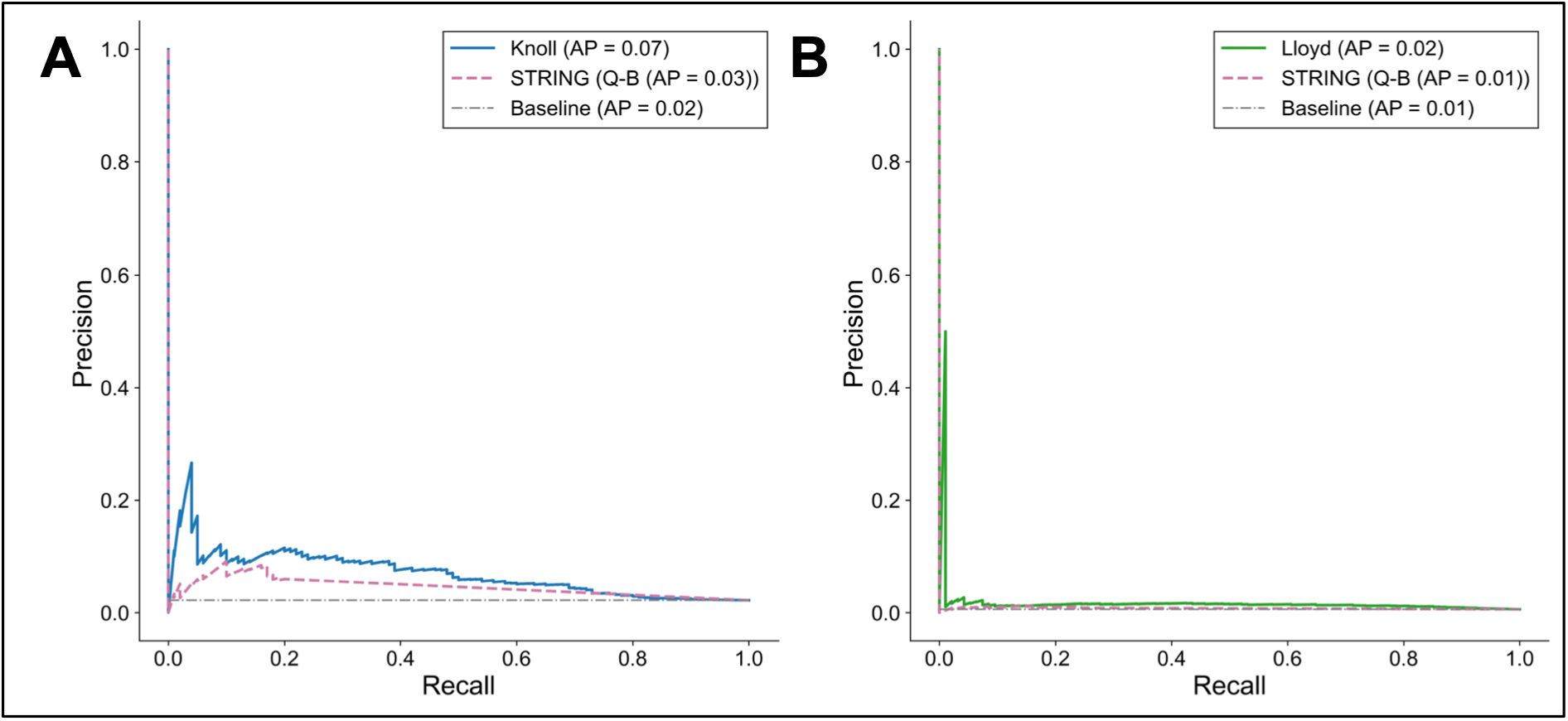
Precision-Recall curves for independent validation screens. **A)** Precision-Recall curve for the MTAP-PRMT5 screen (Knoll et al., 2025). Random forest predictions (blue) were taken directly from the genome-wide MTAP-PRMT5 prediction in our database and matched to the screen genes. STRING interaction scores (pink dashed) and the class-prevalence baseline (grey dash-dotted) are shown for reference. **B)** Precision-Recall curve for the ATM-deficient ATRi screen (Lloyd et al., 2021). random forest predictions (green) were taken from the ATM-ATR prediction, which was generated with the ARID1A-ATR screen excluded to prevent leakage. Baselines as in A. AP, average precision.

